# Effects of Chronic Intermittent Hypoxia and Chronic Sleep Fragmentation on Gut Microbiome, Serum Metabolome, Liver and Adipose Tissue Morphology

**DOI:** 10.1101/2021.11.15.468769

**Authors:** Fan Wang, Juanjuan Zou, Huajun Xu, Weijun Huang, Xiaoman Zhang, Zhicheng Wei, Xinyi Li, Yupu Liu, Jianyin Zou, Feng Liu, Huaming Zhu, Hongliang Yi, Jian Guan, Shankai Yin

## Abstract

Chronic intermittent hypoxia (CIH) and chronic sleep fragmentation (CSF) are two cardinal pathological features of obstructive sleep apnea (OSA). Dietary obesity is a crucial risk intermediator for OSA and metabolic disorders. Gut microbiota affect hepatic and adipose tissue morphology under conditions of CIH or CSF through downstream metabolites. However, the exact relationship is unclear. Herein, chow and high-fat diet (HFD)-fed mice were subjected to CIH or CSF for 10 weeks each and compared to normoxia (NM) or normal sleep (NS) controls. 16S rRNA amplicon sequencing, untargeted liquid chromatography-tandem mass spectrometry, and histological assessment of liver and adipose tissues were used to investigate the correlations between the microbiome, metabolome, and lipid metabolism under CIH or CSF condition. Our results demonstrated that CIH and CSF regulate the abundance of intestinal microbes (such as *Akkermansia mucinphila*, *Clostridium spp.*, *Lactococcus spp.*, *and Bifidobacterium spp*.) and functional metabolites, such as tryptophan, free fatty acids, branched amino acids, and bile acids, which influence adipose tissue and hepatic lipid metabolism, and the level of lipid deposition in tissues and peripheral blood. In conclusion, CIH and CSF adversely affect fecal microbiota composition and function, and host metabolism; these findings provide new insight into the independent and synergistic effects of CIH, CSF, and HFD on lipid disorders.

## 1 Introduction

Obstructive sleep apnea (OSA) is a common sleep-disordered breathing disease that affects almost 1 billion adults worldwide (1). OSA is associated with high economic and health burdens due to the increased risk of cardiovascular and metabolic diseases (2). Exploring the mechanisms underlying the effects of OSA may lead to improved diagnosis and treatment of the disease and its comorbidities. OSA is characterized by recurrent episodes of airway collapse during sleep that cause chronic intermittent hypoxia (CIH) and sleep disruption due to recurrent arousals, called chronic sleep fragmentation (CSF) (3). CIH and CSF are associated with a wide range of neural, hormonal, thrombotic, and metabolic alterations that promoted OSA-related complications (4–6).

In our previous systemic review, we summarized the changes in the metabolome and microbiota that may play an integral role as intermediary factors in the pathophysiology of OSA and related cardiovascular, metabolic, and neurological complications (7). In rodent studies, CIH and CSF induce dysbiosis of the gut microbial community and serum metabolomics changes. In particular, CIH-exposed models showed an altered Firmicutes/Bacteroidetes (F/B) ratio, reduced abundance of Proteobacteria and Clostridia, and changes in molecular metabolites (>22%) suggestive of excessive production of reactive oxygen species (ROS) and free fatty acids (FFA) (8–10). In contrast, long-term CSF-induced gut microbiota changes include increased abundances of Lachnospiraceae and decreased abundance of Lactobacillaceae (11). The level of energy metabolism and monoamine hormones were significantly reduced in CSF mice compared to controls (12). However, few previous studies have comprehensively examined the effects of CIH and CSF on gut microbial composition and function, or the association thereof with host metabolism.

In the present study, we explored the effects of CIH and CSF, with or without dietary intervention, on gut microbial composition, metabolic function, and adipose tissue morphology. We aimed to determine how, and to what extent, CIH and CSF affect hepatic and adipose tissue morphology; changes in gut microbiome and serum metabolites associated with CIH and CSF; and the impact of CIH- and CSF-mediated taxonomic and molecular alterations on hepatic and adipose lipid metabolism.

## 2 Materials and methods

### 2.1 Animals fed a high-Fat Diet

The study protocol was approved by the Institutional Animal Care and Use Committee of Shanghai Jiao Tong University Affiliated Sixth People’s Hospital, China. All experiments were conducted following the Guide for the Care and Use of Laboratory Animals published by the Animal Welfare Committee of the Agricultural Research Organization. Seventy-two male C57BL/6J mice (6 weeks old; Shanghai Laboratory Animal Center, Shanghai, China) were housed in 16 cages (4–5 mice/cage) under standard conditions (temperature: 21 ± 1°C; relative humidity: 50 ± 10%) with a regular 14-h light: 10-h dark cycle (lights on at 7:00 am). After adaptation for 2 weeks, mice (8 weeks old) were randomly divided into eight groups: normoxia (NM; n = 8), CIH (n = 10), HFD (n = 8), CIH+HFD (n = 10), normal Sleep (NS; n = 9), CSF (n = 9), NS+HFD (n = 9) and CSF+HFD (n = 9) groups (Fig. 1A). The mice had free access to food and water for 10 weeks. The chow diet group was fed a low-calorie diet (10% calories from fat), whereas the HFD group was fed a eutrophic diet with 59% calories derived from coconut oil.

**Figure 1.**
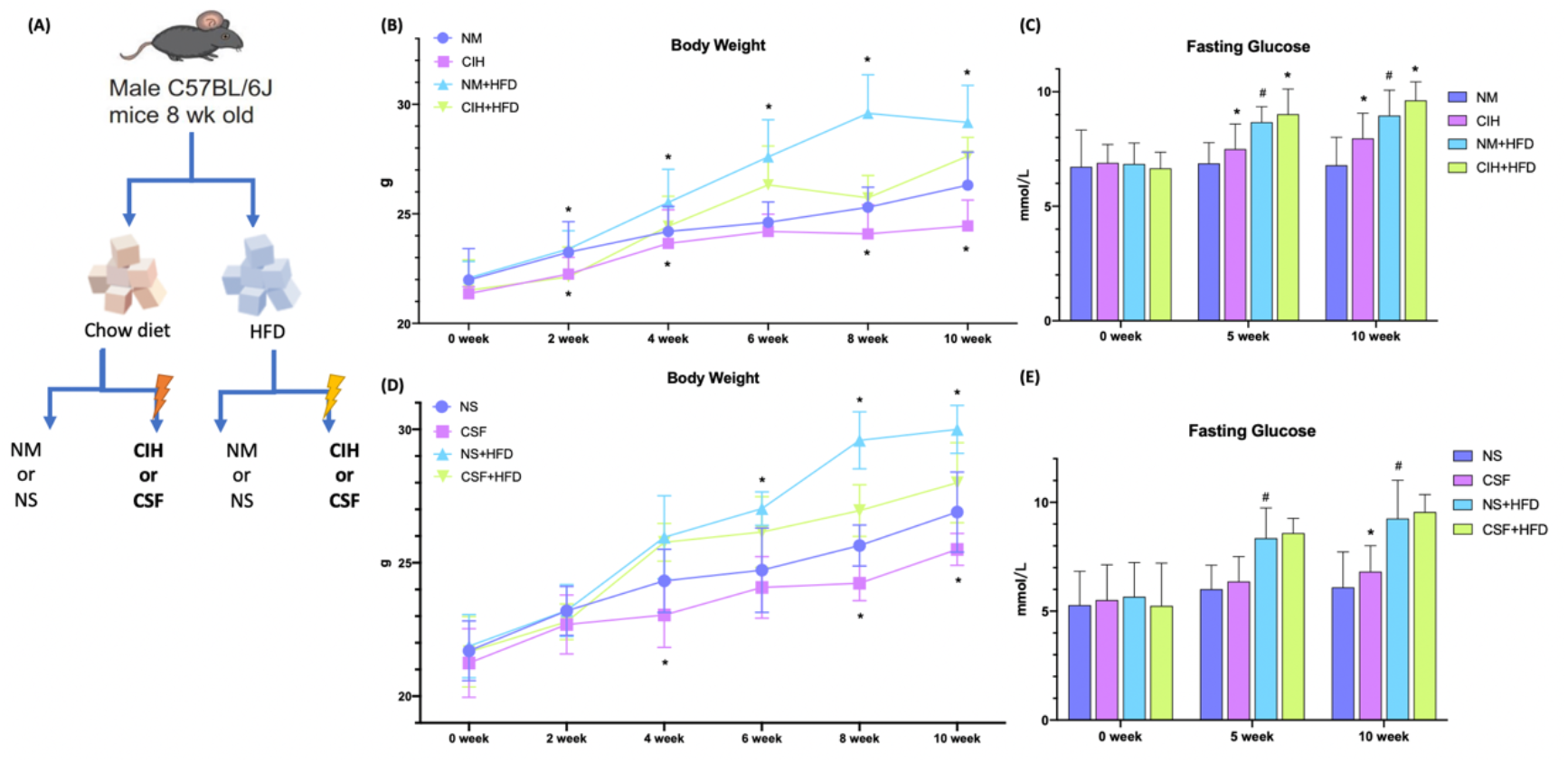
Overall modeling workflow and baseline data. (A) CIH and CSF with diet intervention modeling workflow and design. (B) Body weight of the chow diet and HFD groups throughout the experiment of CIH exposure. (C) The fasting blood glucose levels of the mice in the chow diet and HFD groups during the period of CIH. (D) Body weight of the chow diet and HFD groups throughout the experiment of CSF exposure. (E) The fasting blood glucose levels of the mice in the chow diet and HFD groups during the period of CSF. Data are expressed as means ± SEM. Group differences were assessed by using the Mann-Whitney U test. *P < 0.05 when comparing NM/NS and CIH/CSF groups (NM/NS group vs CIH/CSF group, NM+HFD/NS+HFD group vs CIH+HFD/CSF+HFD group). *^#^*P < 0.05 when comparing chow diet and HFD groups (NM/NS group vs NM+HFD/NS+HFD group).

### 2.2 CIH Exposure

CIH was induced in identical chambers (Oxycycler model A84; BioSpherix, Redfield, NY, USA) for 8h/day, from 8:00 am to 4:00 pm, for 10 weeks. The O_2_ concentration in the chamber was continuously measured using an O_2_ analyzer. Hypoxia was induced with 10% O2 for approximately 240 s, followed by 21% O2 for 120 s, as previously described (13). For NM controls, 10% O2 (hypoxia) was replaced by 21% O2 (air), while the other conditions were similar to those for the CIH groups.

### 2.3 CSF Intervention

Continuous CSF was induced for 10 weeks by using big large, modified, multiple cages equipped with a fixed platform and twirling pole (MaxQ 2000; ThermoFisher Scientific, Waltham, MA, USA), controlled using a timer programmed for random rotation (H3CR-F8-300; Omron Corp., Kyoto, Japan). CSF was induced by rotating the pole for 10 s after every 110 s at a speed of 110 rpm for the entire day to disturb sleeping mice (14).

### 2.4 Hepatic and adipose tissue morphology

The body weight of all mice was recorded weekly and blood glucose (BG) levels were measured every 5 weeks after fasting for 12 h using a glucose analyzer (Accu-Chek; Roche, Rotkreuz Switzerland) over the tail vein. After the 10-week exposure period and 12-h solid and liquid fast, the mice were sacrificed via an overdose of sodium pentobarbital (100 mg/kg, i.p.). Then, samples from the liver, inguinal white adipose tissue (IWAT), perirenal white adipose tissue (PWAT), and scapula brown adipose tissue (BAT) were harvested and weighed. Liver tissues were embedded in Tissue-Tek OCT Compound (Sakura Finetek, Torrance, CA, USA), sliced into cryosections (8-μm-thick) and stained with Oil Red O (Sigma-Aldrich, St. Louis, MO, USA) to assess the fatty droplet content. The stained liver cells were analyzed using Image Pro-Plus 6.0 (Media Cybernetics, Bethesda, MD, USA). Adipose tissues were fixed in 10% formalin, embedded in paraffin wax, sectioned (5 μm), and stained with hematoxylin and eosin. Hematoxylin and eosin staining was used to evaluate the sizes of subcutaneous and visceral white adipose tissue (WAT) and BAT. Mean cell size and size distribution was determined for WAT. Because it was difficult to isolate the BAT cells due to cytoplasmic lipid accumulation, the lipid area of BAT was quantified. The lipid area and cell size were calculated using MetaMorph imaging software (Molecular Devices, Downingtown, PA, USA).

### 2.5 Microbiome assessment

Fecal samples were collected from all groups for microbiota 16S rRNA analysis. The microflora were detected by Lianchuan Biotechnology Co., Ltd. (Hangzhou, China). Bacterial genomic DNA was obtained from frozen colon contents using the QIAamp DNA Stool Mini Kit (51504; Qiagen, Germantown, MD, USA), according to the manufacturer’s instructions. Successful DNA isolation was confirmed by agarose gel electrophoresis. DNA samples were amplified by polymerase chain reaction (PCR) using bar-coded primers flanking the V3–V4 region of the 16S rRNA gene. The PCR amplicon products were separated on 0.8% agarose gels and extracted. Only PCR products without primer dimers and contaminant bands were used for sequencing by synthesis. High-throughput pyrosequencing of the PCR products was performed using the Illumina MiSeq platform (Illumina, Inc., San Diego, CA, USA). Paired-end reads of the original DNA fragments were excluded from the analysis if they did not well-match a 12-base Golay barcode (one or no errors), the read overlap was < 35 bases, the overlapped region differed by more than 15%, or there were more than three base calls below Q20. Bacterial operational taxonomic units (OTUs) were created by clustering the reads at 97% identity in the Quantitative Insights into Microbial Ecology (QIIME; http://qiime.org/scripts/pick_otus.html) database.

### 2.6 Metabolome assessment

Venous blood samples were obtained from mice after overnight fasting. Peripheral blood was allowed to clot for 30 min at room temperature and centrifuged at 3,000 rpm for 10 min to collect the supernatant. Serum samples were immediately stored as aliquots at −80℃ prior to further sample preparation and analysis. A total of 50 µL of serum from each of the eight groups was used for metabolic profiling. Untargeted metabolites were detected by Lianchuan Biotechnology. Briefly, serum metabolites were measured by direct-injection electrospray tandem mass spectrometry (MS/MS) using a Micromass Quattro Micro liquid chromatography (LC)-MS system (Waters-Micromass, Milford, MA, USA) equipped with a model HTS-PAL 2777 autosampler (Leap Technologies, Carrboro, NC, USA), model 1525 high-performance liquid chromatography (HPLC) solvent delivery system (Agilent Technologies, Palo Alto, CA, USA), and MassLynx data system (version 4.0; Waters-Micromass). For each batch, periodic analysis of the same pooled quality control (QC) sample was performed to detect variations within and between experiments (15). The supplementary tables summarize the amino acids, organic carbohydrate acids, and lipid-related molecules.

### 2.7 Statistical analysis

Taxonomy abundance was normalized to the summed OTUs of each sample. Principal coordinate analysis (PCoA) of β-diversity was performed; the (unpaired) Wilcoxon rank-sum test was used to identify the most diverse taxa. Phylogenetic Investigation of Communities by Reconstruction of Unobserved States (PICRUSt) analysis was applied to metagenome functions predicted from the normalized OTUs for Kyoto Encyclopedia of Genes and Genomes (KEGG) orthologs (16). The Mann-Whitney U test and Spearman’s correlation were performed using SPSS software (version 26.0; IBM Corp., Armonk, NY, USA). Data are expressed as mean ± SEM. Significance was set at p < 0.05* or p < 0.01**.

## 3 Results

### 3.1 Effects of CIH and CSF on body, liver, and adipose tissue weight, and glucose level

All mice survived and were kept in good condition during the 10-week-long CIH or CSF exposure with or without a HFD (Fig. 1A). The body weight significantly decreased after 2 weeks under CIH in both the chow- and HFD-fed groups compared to the control groups (Fig. 1B). Similarly, a 4-week-long CSF intervention down-regulated the weight gain in mice irrespective of the diet (Fig. 1D). The body weight was significantly higher in the NM and NS+HFD groups than the other groups (Fig. 1B, D). In later stages, the change in body weight in the eight groups decreased. Fasting BG level increased with CIH and HFD interventions, and was higher in the CIH+HFD group compared to the other groups (Fig. 1C). The fasting BG level was significantly higher in the NS+HFD compared to CSF group (Fig. 1E).

Red oil staining of the liver tissues (Fig. 2B) revealed that 10-week CIH exposure sharply increased microbubble lipid droplets in liver cells, particularly around the hepatic sinus, compared to the NM group. HFD-fed mice showed marked homogeneous steatosis in parenchymal hepatocytes. The CIH+HFD group demonstrated excessive deposition of microbubble and diffuse bullae lipid droplets, disordered hepatic cords, and higher liver weight compared to the control group (Fig. 2A, B).

**Figure 2.**
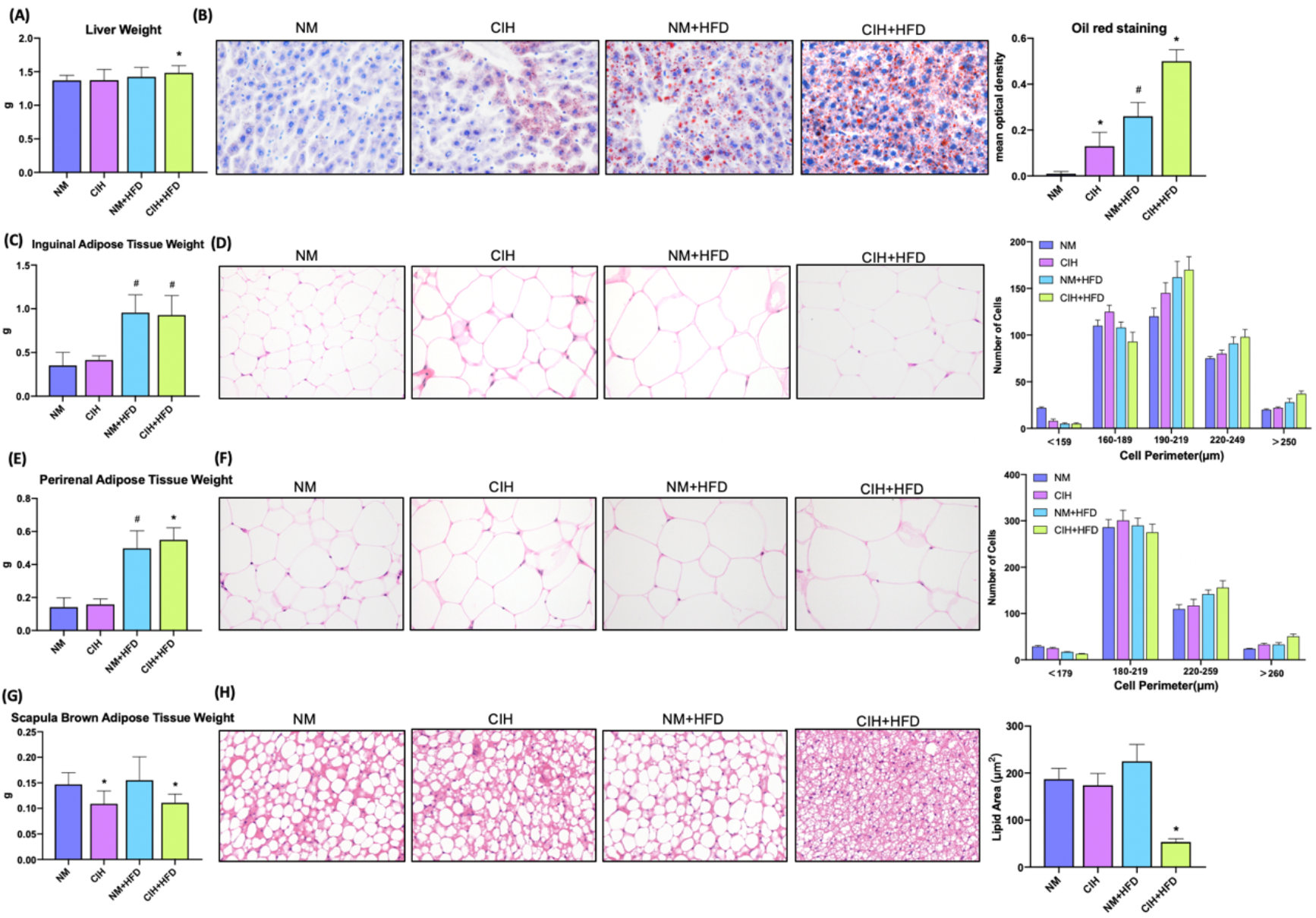
Profiles of liver tissue, abdominal subcutaneous and visceral white adipose tissue, and scapula brown adipose tissue of mice exposed to CIH intervention. (A) Liver weight of the mice in the chow diet and HFD groups pre- and post- CIH. (B) Oil red staining of liver tissue and mean optical density of the mice in the chow diet and HFD groups pre- and post- CIH. (C) Inguinal adipose tissue (Abdominal subcutaneous white adipose tissue) weight of the mice in the normal and HFD groups before and after CIH. (D) Hematoxylin-eosin staining and the distribution of adipocyte perimeters of inguinal adipose tissue of the chow diet-fed and HFD-fed mice before and after CIH. (E) Perirenal adipose tissue (abdominal visceral white adipose tissue) weight of the mice in the chow diet and HFD groups pre- and post- CIH. (F) Hematoxylin-eosin staining and the distribution of adipocyte perimeters of perirenal adipose tissue of the mice and brown fat weight of the chow diet-fed and HFD-fed mice before and after CIH. (G) Scapula brown adipose tissue weight of the mice in the chow diet and HFD groups pre- and post- CIH. (H) Hematoxylin-eosin staining and lipid area of scapula brown adipose tissue of the mice in the chow diet and HFD groups pre- and post- CIH. Data are expressed as means ± SEM. Group differences were assessed by using the Mann-Whitney U test. *P < 0.05 when comparing NM and CIH groups (NM group vs CIH group, NM+HFD group vs CIH+HFD group). *^#^*P < 0.05 when comparing chow diet and HFD groups (NM group vs NM+HFD group). Original magnification: 400 ×.

Liver weight (Fig. 3A) markedly decreased after 10 weeks of CSF with or without HFD. In contrast to the CIH group, the CSF group did not demonstrate increased deposition of microbubble lipid droplets in liver cells (Fig. 3B). The liver morphology of CSF+HFD mice was similar to that of HFD mice, except for hepatic steatosis. Moreover, tissue distant from the hepatic sinuses showed vacuolar degeneration and sporadic nucleolysis in the CSF+HFD group.

**Figure 3.**
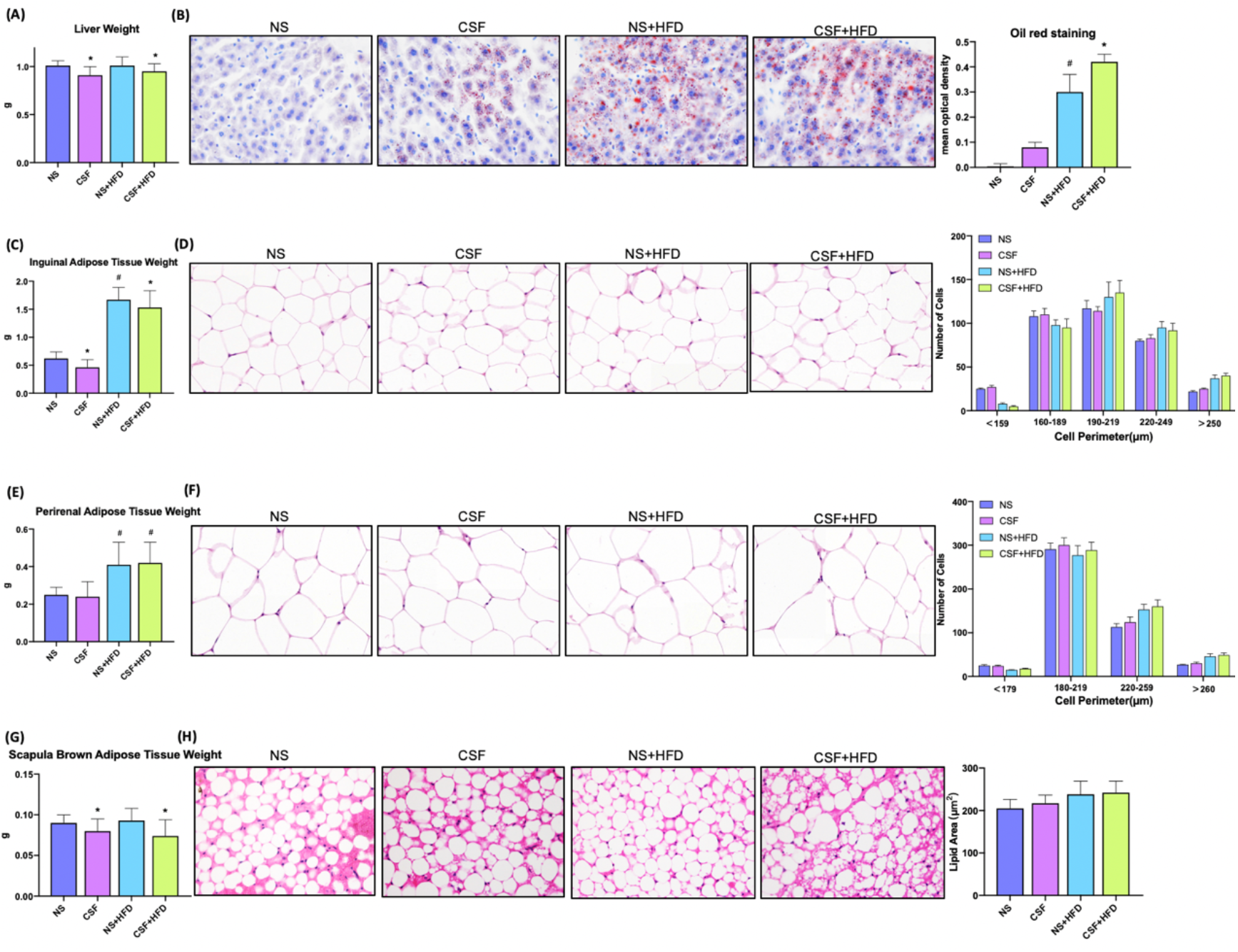
Profiles of liver tissue, abdominal subcutaneous and visceral white adipose tissue, and scapula brown adipose tissue of mice exposed to CSF intervention. (A) Liver weight of the mice in the chow diet and HFD groups pre- and post- CSF. (B) Oil red staining of liver tissue and mean optical density of the mice in the chow diet and HFD groups pre- and post- CSF. (C) Inguinal adipose tissue (Abdominal subcutaneous white adipose tissue) weight of the mice in the normal and HFD groups before and after CSF. (D) Hematoxylin-eosin staining and the distribution of adipocyte perimeters of inguinal adipose tissue of the chow diet-fed and HFD-fed mice before and after CSF. (E) Perirenal adipose tissue (abdominal visceral white adipose tissue) weight of the mice in the chow diet and HFD groups pre- and post- CSF. (F) Hematoxylin-eosin staining and the distribution of adipocyte perimeters of perirenal adipose tissue of the mice and brown fat weight of the chow diet-fed and HFD-fed mice before and after CSF. (G) Scapula brown adipose tissue weight of the mice in the chow diet and HFD groups pre- and post- CSF. (H) Hematoxylin-eosin staining and lipid area of scapula brown adipose tissue of the mice in the chow diet and HFD groups pre- and post- CSF. Data are expressed as means ± SEM. Group differences were assessed by using the Mann-Whitney U test. *P < 0.05 when comparing NS and CSF groups (NS group vs CSF group, NS+HFD group vs CSF+HFD group). #P < 0.05 when comparing chow diet and HFD groups (NS group vs NS+HFD group). Original magnification: 400 ×.\

The weights of abdominal subcutaneous and visceral WAT were similar in all chow-fed mice, irrespective of CIH stimulation (Fig. 2C, E). Hematoxylin and eosin staining showed enlarged lipid droplets in white adipocytes in the CIH mice (Fig. 2D, F). The WAT cells were vacuolated and disorganized in HFD-fed mice. The PWAT weight was significantly higher in the CIH+HFD compared to HFD mice (Fig. 2E). The cytoderm of hypertrophic IWAT in the CIH+HFD group was thin and irregularly shaped (Fig. 2D).

CSF intervention was associated with reduced IWAT weight (Fig. 3C), but not morphological variation of adipose cells (Fig. 3D). The PWAT weight appeared to be influenced by the HFD diet, but not by CSF exposure (Fig. 3E). The majority of WAT cells were transformed from smaller adipose cells to loose cells in the HFD and CSF+HFD groups compared to the controls (Fig. 3D, F).

Weight of the scapula BAT significantly decreased in the CIH and CSF groups despite HFD (Figs. 2G and 3G). Fibrosis and lipid fragmentation were observed in the BAT due to the combined effects of HFD and CIH or CSF stimulation (Figs. 2H and 3H).

### 3.2 Effects of CIH and CSF on fecal microbiota

#### 3.2.1 Effects of CIH on fecal microbiota

The PCoA plot demonstrated that the NM and CIH groups were separated along the PC1 axis, whereas the NM+HFD and CIH+HFD groups were separated along the PC2 axis (Fig. 4A). We further evaluated group similarities at the phylum level via hierarchical clustering of the four groups (Fig. 4B). CIH significantly changed the microbial composition in both chow- and HFD-fed mice. All mice had similar microbiome compositions after CIH exposure, irrespective of their dietary and metabolic patterns at baseline. Analysis of phylum abundance revealed that the predominant bacteria in the cecum were Firmicutes, Bacteroidetes, Actinobacteria, Proteobacteria, and Patescibacteria (Fig. 4C). Moreover, CIH significantly increased the abundance of Firmicutes and Proteobacteria, and decreased that of Bacteroidetes in both chow- and HFD-fed mice. The abundance of Actinobacteria was increased in the NM+HFD and CIH+HFD groups as a result of the change in dietary pattern or additive effect of CIH.

**Figure 4.**
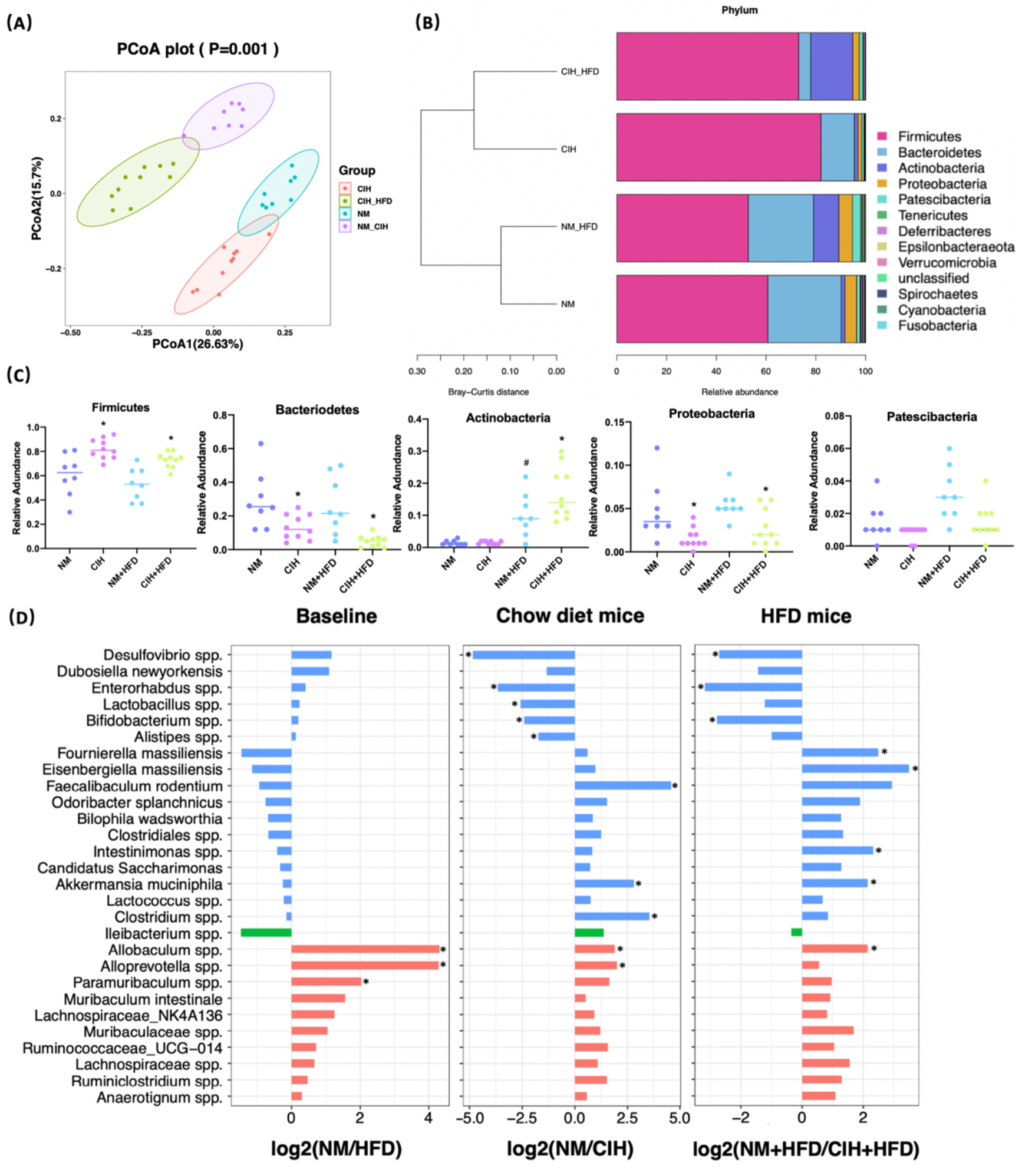
Effects of CIH on fecal microbiota. (A) PCOA plot generated by using microbiota OTU metrics based on the Bray-Curtis similarity for the NM, CIH, NM+HFD, CIH+HFD groups (n = 8-10/group). (B) The hierarchical cluster based on the Bray-Curtis similarity of the samples from the NM, CIH, NM+HFD, CIH+HFD groups. The bar plot shows the abundance of each phylum in each sample. (C) The 5 most abundant phyla in the NM, CIH, NM+HFD, CIH+HFD groups. Data are expressed as means ± SEM. Group differences were assessed by using the Mann-Whitney U test. *P < 0.05 when comparing NM vs CIH, or NM+HFD vs CIH+HFD, #P < 0.05 when comparing NM vs NM+HFD. D) Fold changes of the annotated microbes at the genus level. The data of log2(fold change) are expressed as means. Group differences were assessed by using the Mann-Whitney U test. *P < 0.05.

We further evaluated differences in the cecal microbial species at the genus level. Twenty-eight microbes were selected, as there was no “0” relative abundance value in any CIH model. Figure 4D illustrates the log2 fold change (log2FC) in the NM/NM+HFD (baseline difference), NM/CIH (CIH-induced changes in chow-fed mice), and NM+HFD/CIH+HFD (CIH-induced changes in HFD mice) groups. Red bars indicate the same direction of log2FC change in the three NM/CIH and NM+HFD/CIH+HFD pairwise group comparisons. The highest log2FC value was observed for *Allobaculum spp.* Green bars indicate the same direction of log2FC change for the NM/NM+HFD and NM+HFD/CIH+HFD groups; the changes in *Ileibacterium spp.* were primarily induced by dietary patterns. Blue bars indicate microbes with the same direction of log2FC change for the NM/CIH and NM+HFD/CIH+HFD groups, suggesting that the variation may have been induced by CIH. Importantly, CIH significantly increased the abundances of *Desulfovibrio spp.*, *Enterohabdus spp.*, and *Bifidobacterium spp.* in chow- and HFD-fed mice (log2FC < 0; p < 0.05). CIH alone significantly increased the abundances of *Lactobacillus spp. and Alistipes spp.*, while decreasing the abundances of *Feacalibaculum rodentium* and *Clostridium spp.* (log2FC > 0; p < 0.05). The CIH+HFD intervention down-regulated the abundances of *Fournierella massiliensis*, *Eisenbergiella massiliensis*, *Intestinimonas spp.*, and *A. muciniphila*.

#### 3.2.2 Effects of CSF on fecal microbiota

The PCoA plot (Fig. 5A) showed that the NS and CSF groups were partially separated along the PC1 axis, whereas the NS+HFD and CSF+HFD groups were fully separated along the PC2 axis. “Stacked clustering” (Fig. 5B) of the four groups revealed that CSF exposure slightly altered the microbial composition of chow-fed mice compared to HFD-fed mice. In contrast, the microbial composition was significantly changed after the combined CSF+HFD intervention.

**Figure 5.**
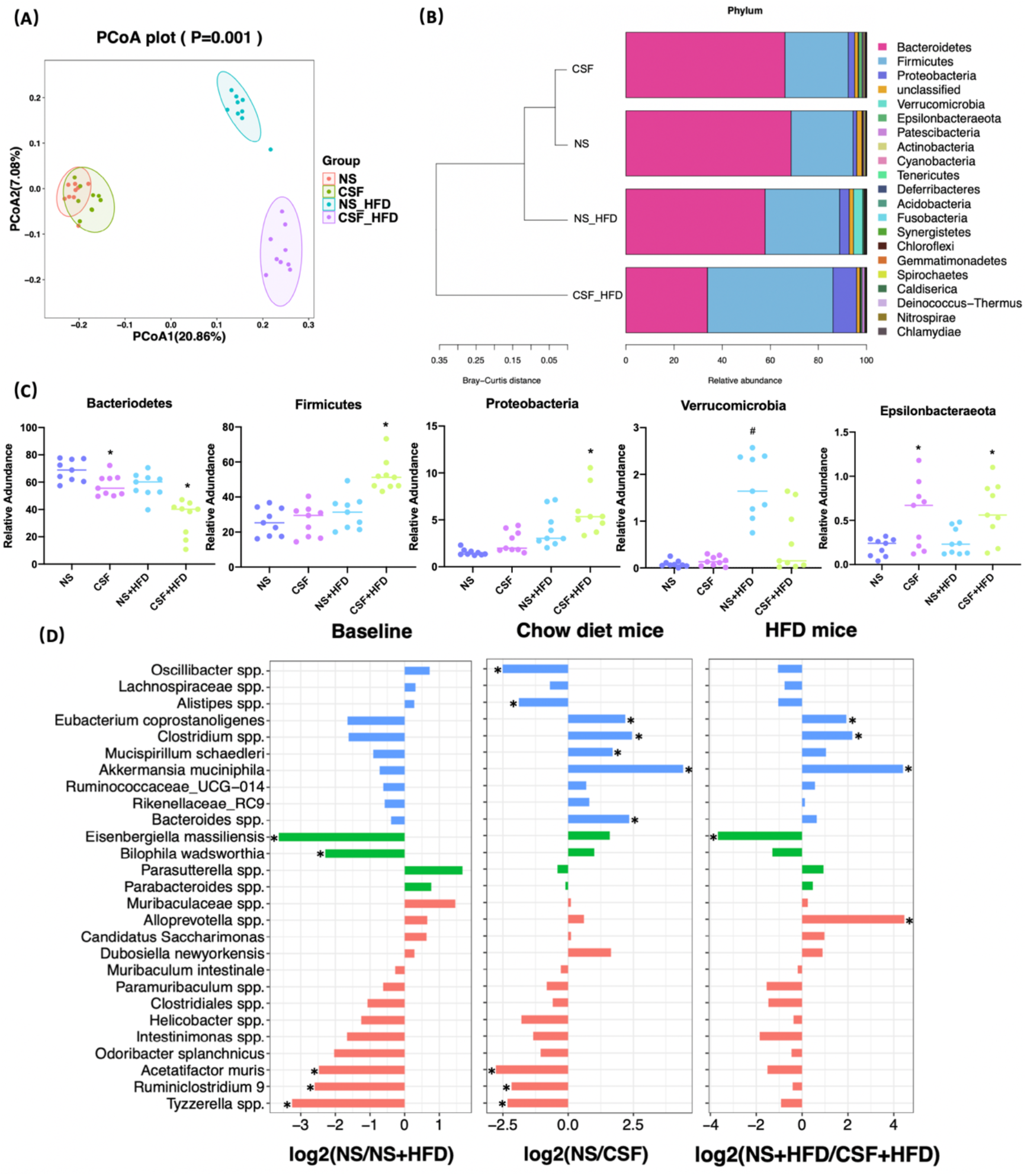
Effects of CSF on fecal microbiota. (A) PCOA plot generated by using microbiota OTU metrics based on the Bray-Curtis similarity for the NS, CSF, NS+HFD, CSF+HFD groups (n = 9/group). (B) The hierarchical cluster based on the Bray-Curtis similarity of the samples from the NS, CSF, NS+HFD, CSF+HFD groups. The bar plot shows the abundance of each phylum in each sample. (C) The 5 most abundant phyla in the NS, CSF, NS+HFD, CSF+HFD groups. Data are expressed as means ± SEM. Group differences were assessed by using the Mann-Whitney U test. *P < 0.05 when comparing NS vs CSF, or NS+HFD vs CSF+HFD, #P < 0.05 when comparing NS vs NS+HFD. D) Fold changes of the annotated microbes at the genus level. The data of log2(fold change) are expressed as means. Group differences were assessed by using the Mann-Whitney U test. *P < 0.05.

Analysis of the phyla of microbiota demonstrated that Bacteroidetes, Firmicutes, Proteobacteria, Verrucomicrobia, and Epsilonbacteraeota dominated the fecal bacterial population (Fig. 5C). CSF exposure significantly decreased the abundance of Bacteroidetes, but increased the abundances of Firmicutes and Proteobacteria in HFD-fed mice. Irrespective of the dietary pattern, the abundance of Epsilonbacteraeota increased after CSF.

Twenty-seven species labeled at the genus level were selected to compare the log2FC (Fig. 5D) between groups. CSF significantly reduced the abundances of *Eubacterium coprostanoligenes*, *Clostridium spp.*, and *A. muciniphila*, irrespective of dietary intervention. CSF exposure alone significantly upregulated *Oscillibacter spp.* and *Alistipes spp.*, and downregulated *Mucispirillum schaedleri* and *Bacteroides spp*.

### 3.3 Effects of CIH and CSF on serum metabolome

#### 3.3.1 Effect of CIH on serum metabolome

We detected 16,493 peaks in the positive and negative ionization modes, likely induced by CIH. After excluding the natural isotopic peaks, 320 and 516 metabolites were identified in the positive and negative modes, respectively, including amino acids, carbohydrates, lipid molecules, bile acids (BA), fatty acids and vitamins. Unsupervised principal component analysis (PCA) analysis was used to evaluate the intrinsic metabolic variations. Metabolites from the CIH group were more separated than those from the NM control group. However, metabolites from the CIH+HFD group showed less obvious separation than those from the NM+HFD group, indicating more significant metabolic differences between the CIH and NM groups (Fig. 6A). Tight clustering of the QC samples indicated good reproducibility across samples. Volcano plots (Fig. 6B) were constructed to evaluate the differential metabolites between the NM and CIH groups, and between the NM+HFD and CIH+HFD groups. The statistical significance level in the clustering analysis was a false discovery rate < 0.05 (> 4/3- or < 3/4-fold change).

**Figure 6.**
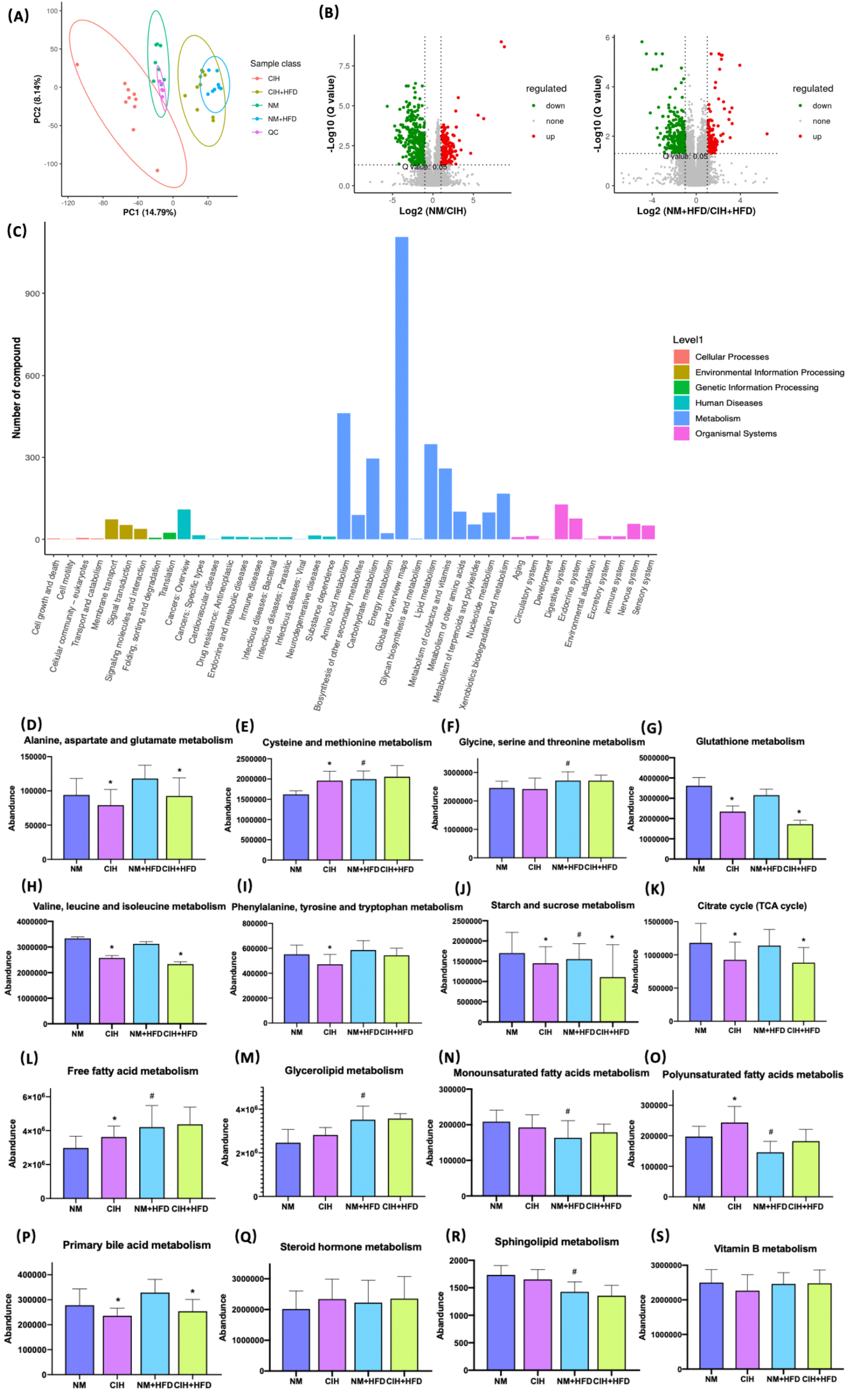
Effects of CIH on the serum metabolome. (A) The PCA plot of metabolites generated by using the metabolite intensity from the NM (n = 8), CIH (n = 10), NM+HFD (n = 8), and CIH+HFD (n = 10) groups. (B) Volcano plots of metabolites with FDR < 0.05 and fold changes of > 4/3 or < 3/4 when comparing NM vs CIH, or comparing NM+HFD vs CIH+HFD. (C) Enrichment of activated pathways induced by alterations of metabolites in NM, CIH, NM+HFD and CIH+HFD groups. (D-S) The abundance of metabolite pathways from the NM, CIH, NM+HFD and CIH+HFD groups. Data are expressed as means ± SEM. Group differences were assessed by using the Mann-Whitney U test. *P < 0.05 when comparing NM vs CIH, or NM+HFD vs CIH+HFD, *^#^*P < 0.05 when comparing NM vs NM+HFD.

Figure 6C shows the enrichment of activated pathways, focusing on metabolism-related pathways. We further analyzed the effects of CIH on host metabolic alterations in terms of the abundance of specific metabolic pathways. The abundance of a metabolic pathway from a single sample was given by the sum of the abundance of the metabolites typically found in that pathway (see supplementary tables). Figure 6D–I shows that CIH inhibited the metabolism of alanine, aspartate, glutamate, glutathione, valine, leucine, and isoleucine (i.e., branched-chain amino acids, BCAAs), despite HFD. CIH or HFD intervention alone promoted cysteine and methionine metabolism. Phenylalanine, tyrosine, and tryptophan metabolism were downregulated by CIH, which was partially reversed by HFD. Figure 6J–K shows that CIH inhibited starch and sucrose metabolism, and the citrate cycle. Additionally, Figure 6L–R shows that both CIH and HFD increased FFA metabolism. The increased metabolism of glycerolipid, monounsaturated fatty acids (MUFAs), and sphingolipids due to CIH was further increased after HFD. CIH significantly increased polyunsaturated fatty acid (PUFA) metabolism but decreased the level of primary BA, opposite to the effects of HFD.

#### 3.3.2 Effect of CSF on serum metabolome

In the CSF group, we identified 7,670 natural isotopic peaks (280 and 405 metabolites in positive and negative ionization modes, respectively). The PCA plot (Fig. 7A) showed that metabolites from the CSF+HFD group were more separated than those from the NS+HFD group, whereas those from the CSF group showed less separation than those from the NS control group, suggesting fewer metabolic differences between the latter two groups.

**Figure 7.**
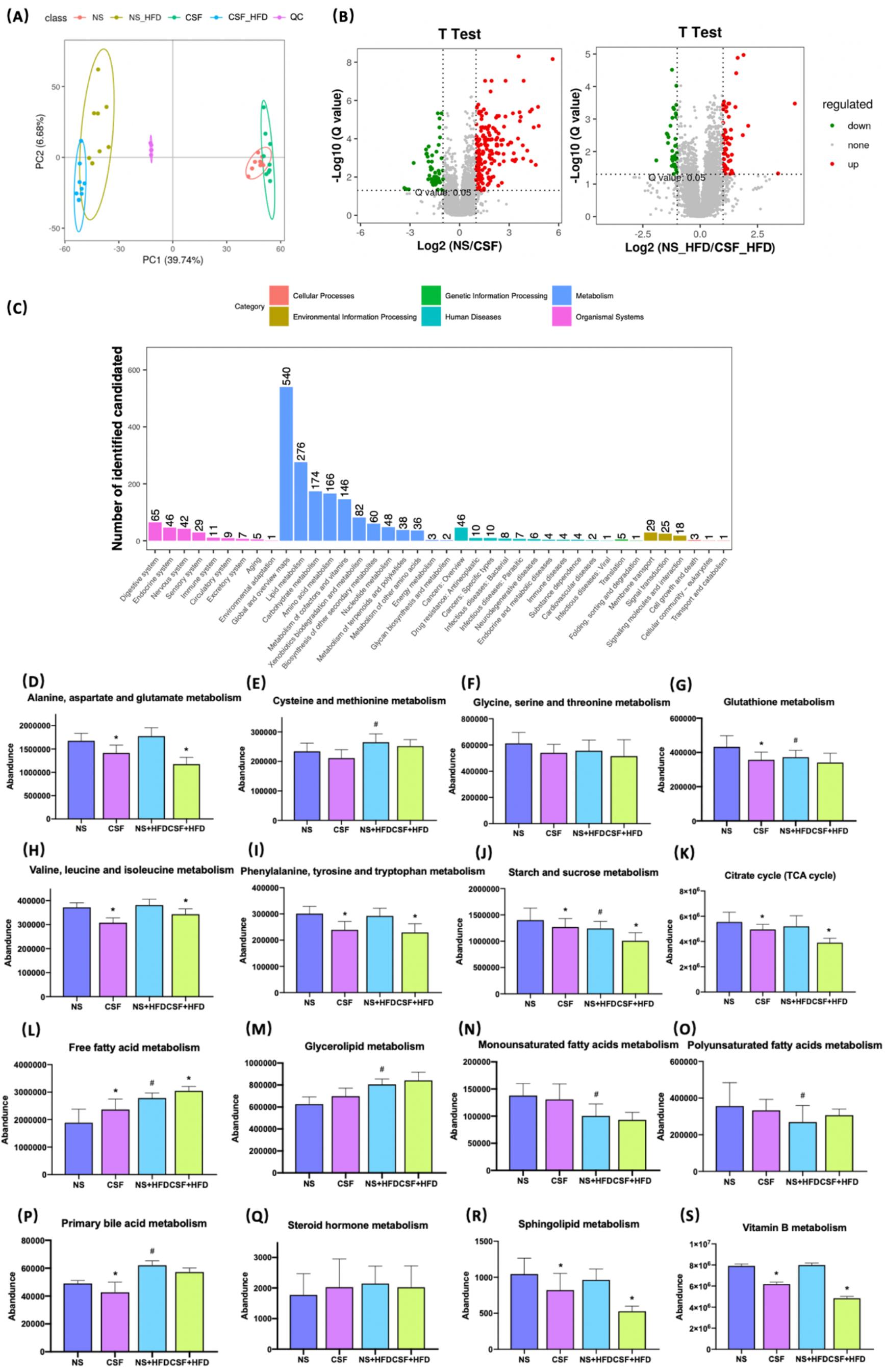
Effects of CSF on the serum metabolome. (A) The PCA plot of metabolites generated by using the metabolite intensity from the NS (n = 9), CSF (n = 9), NS+HFD (n = 9), and CSF+HFD (n = 9) groups. (B) Volcano plots of metabolites with FDR < 0.05 and fold changes of > 4/3 or < 3/4 when comparing NS vs CSF, or comparing NS+HFD vs CSF+HFD. (C) Enrichment of activated pathways induced by alterations of metabolites in NS, CSF, NS+HFD and CSF+HFD groups. (D) The abundance of metabolite pathways from the NS, CSF, NS+HFD and CSF+HFD groups. Data are expressed as means ± SEM. Group differences were assessed by using the Mann-Whitney U test. *P < 0.05 when comparing NS vs CSF, or NS+HFD vs CSF+HFD, *^#^*P < 0.05 when comparing NS vs NS+HFD.

We explored the impact of CSF on host serum metabolic alterations using Volcano plots (Fig. 7B) and enrichment of KEGG metabolic pathways (Fig. 7C). Figure 7D–I showed that CSF inhibited the metabolism of alanine, aspartate, glutamate, glutathione, BCAAs, phenylalanine, tyrosine, and tryptophan. CSF had a synergistic effect with HFD, suppressing starch and sucrose metabolism, and the citrate cycle (Fig. 7J–K). Additionally, CSF significantly increased the level of FFAs and decreased the levels of primary BA and sphingolipids (Fig. 7L–R). Figure 7S shows that vitamin B metabolism was inhibited by CSF.

### 3.4 Effects of CIH and CSF on microbial function

We evaluated the function of gut microbiota using KEGG and PICRUSt analyses. Group differences in microbial function related to the metabolism of amino acids, carbohydrates, lipids, BAs, and FFAs. Figure 8A shows that CIH significantly inhibited BCAA biosynthesis but promoted their degradation, similar to the effects of CSF (Fig. 9A). Similarly, CIH or CSF exposure significantly inhibited alanine, aspartate, glutamate, phenylalanine, tyrosine, tryptophan, and glutathione metabolism, which was attenuated by HFD. Both the CIH and HFD interventions significantly increased the metabolism of cysteine, methionine, and FFAs, whereas they inhibited glutathione metabolism and carbohydrate digestion and absorption. CIH significantly increased the levels of MUFAs and decreased the level of primary BA, opposite to the effects of HFD. CIH in conjunction with HFD predisposed the mice to metabolic and cardiovascular diseases (CVDs). Similarly, CSF reduced the metabolism of glycine, serine, threonine, and primary BAs, and inhibited carbohydrate digestion and absorption. In particular, CSF exposure substantially inhibited the metabolism of sphingolipids and vitamin B, thereby predisposing the mice to neurodegenerative diseases (NDDs) and infectious diseases (IDs).

**Figure 8.**
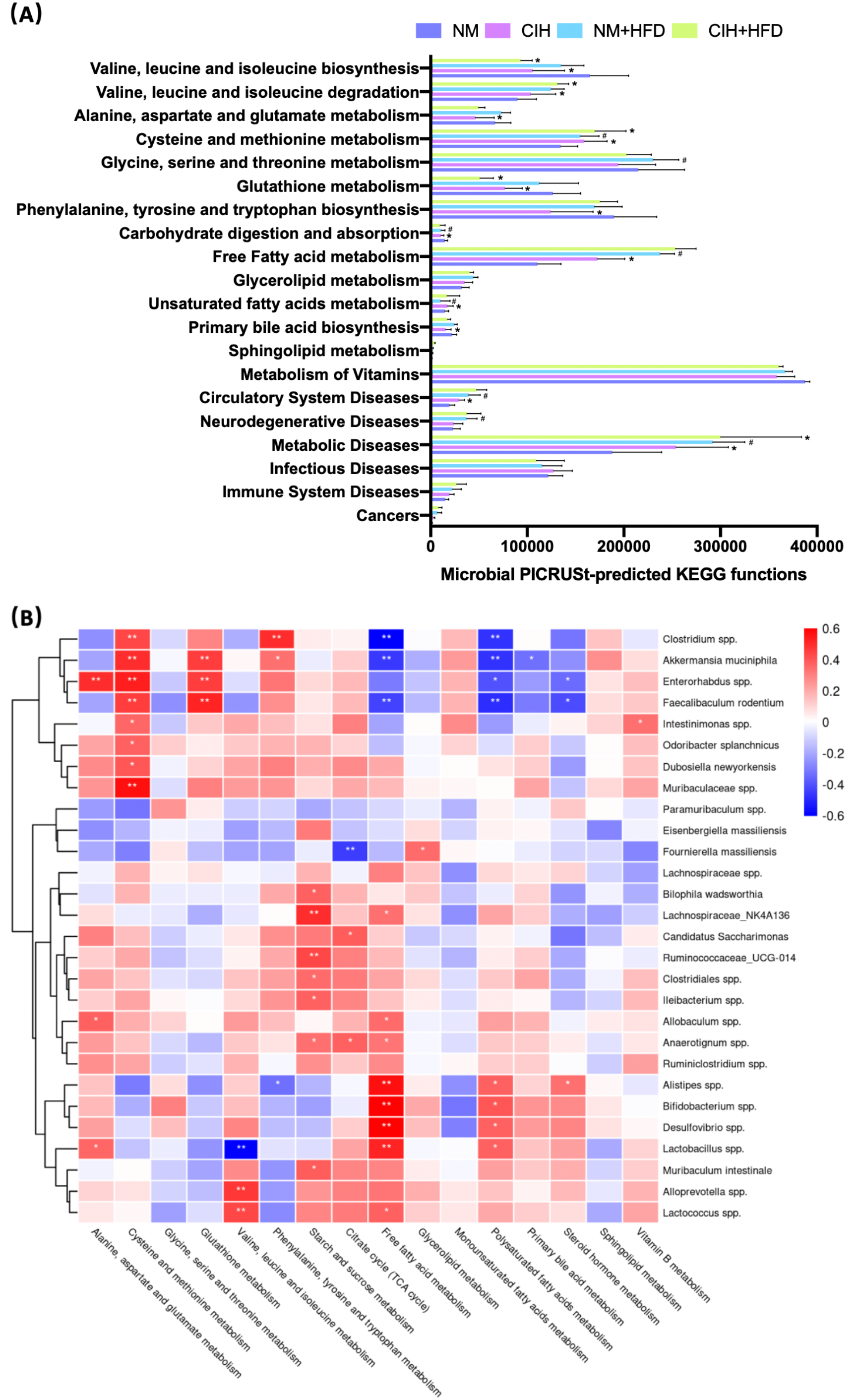
CIH-related fecal microbiota functional changes and correlations between the host metabolome and fecal microbiome. (A) Microbial PICRUSt-predicted KEGG functions relevant to metabolism and diseases. Data are expressed as means ± SEM. Group differences were assessed by using the Mann-Whitney U test. *P < 0.05 when comparing NM vs CIH, or NM+HFD vs CIH+HFD. *^#^*P < 0.05 when comparing NM vs NM+HFD. (B) Spearman correlations of the relative abundance of fecal microbial genus or species and the abundance of metabolite pathways in host serum (n = 36). The r values are represented by gradient colors, with red cells indicating positive correlations and blue cells indicating negative correlations. *P < 0.05, and **P < 0.01.

**Figure 9.**
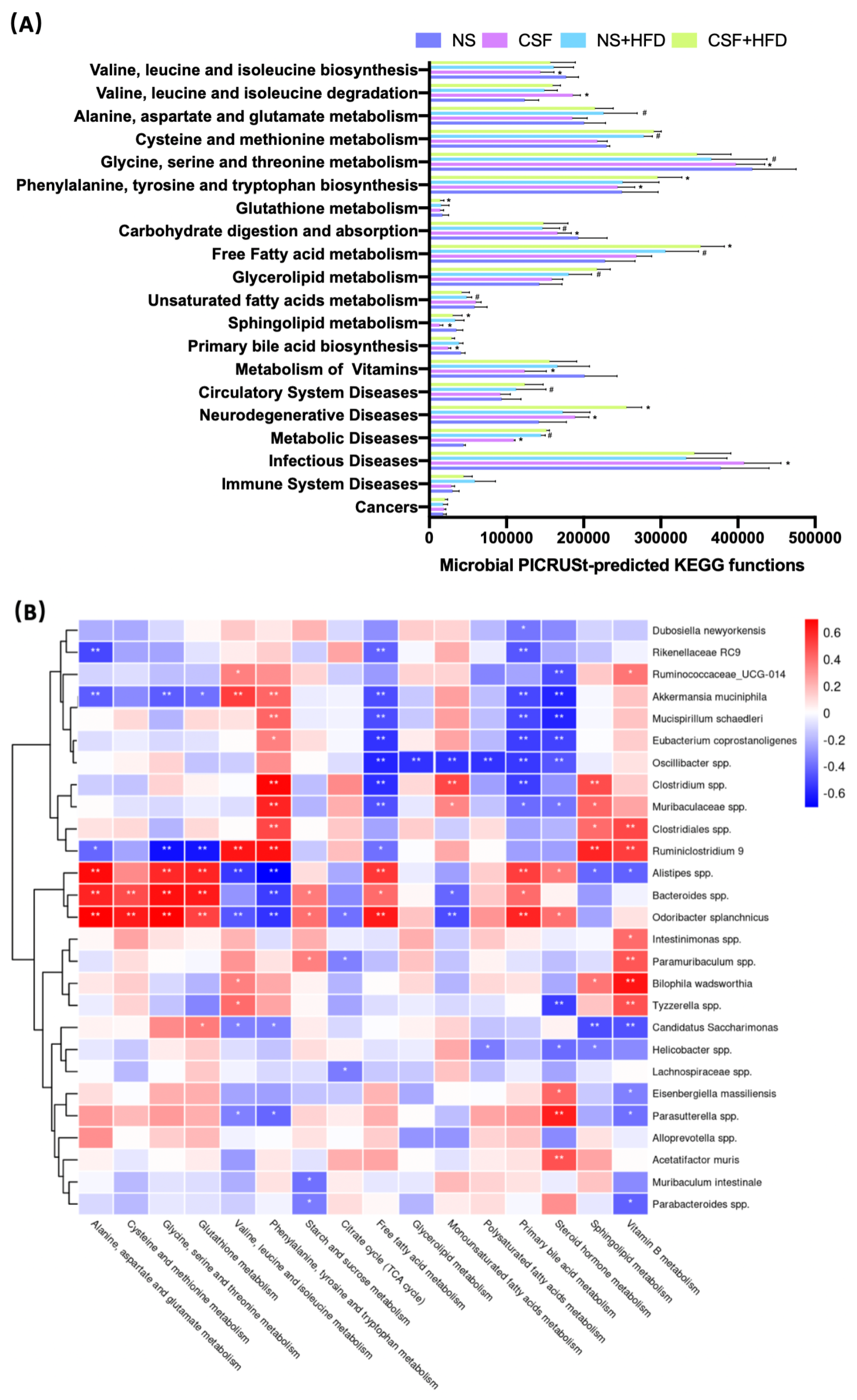
CSF-related fecal microbiota functional changes and correlations between the host metabolome and fecal microbiome. (A) Microbial PICRUSt-predicted KEGG functions relevant to metabolism and diseases. Data are expressed as means ± SEM. Group differences were assessed by using the Mann-Whitney U test. *P < 0.05 when comparing NS vs CSF, or NS+HFD vs CSF+HFD. *^#^*P < 0.05 when comparing NM vs NM+HFD. (B) Spearman correlations of the relative abundance of fecal microbial genus or species and the abundance of metabolite pathways in host serum (n = 36). The r values are represented by gradient colors, with red cells indicating positive correlations and blue cells indicating negative correlations. *P < 0.05, and **P < 0.01.

### 3.5 Correlations of microbiota composition with host metabolism and hepatic and adipose tissue phenotypes

We evaluated the relative abundance of fecal microbes and metabolic pathway activation in CIH host serum and hepatic and adipose tissue phenotypes (Figs. 8B and 10A). Several microbes, such as *A. muciniphila*, *Bifidobacterium spp.*, *Lactobacillus spp.*, and *Clostridium spp.*, were significantly correlated with multiple host metabolic pathways, such as the TCA cycle and metabolism of starch, sucrose, FFA, glutathione, and primary BAs.

**Figure 10.**
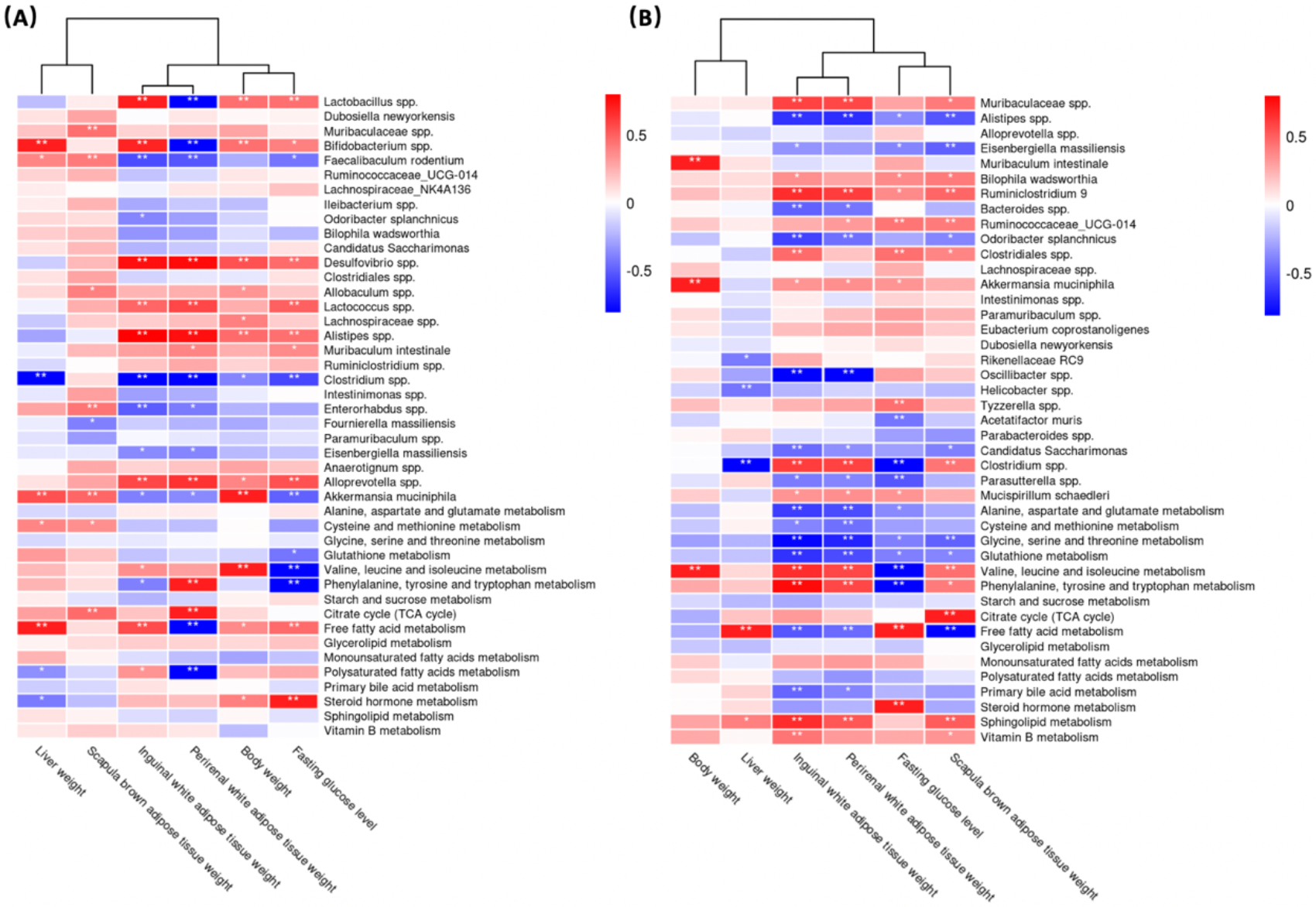
Correlations between the host metabolome, fecal microbiome and hepatic- and adipose-phenotype. (A) Spearman correlations of the relative abundance of fecal microbial genus or species, the abundance of metabolite pathways and phenotype of obesity and fasting glucose in host serum of 4 CIH-related modeling groups (n = 36). (B) Spearman correlations of the relative abundance of fecal microbial genus or species, the abundance of metabolite pathways and phenotype of obesity and fasting glucose in host serum of 4 PSD-related modeling groups (n = 36). The r values are represented by gradient colors, with red cells indicating positive correlations and blue cells indicating negative correlations. *P < 0.05, and **P < 0.01.

CSF stimulation was associated with changes in the abundances of *A. muciniphila*, *M. schaedleri*, *Oscillibacter spp.*, *Bacteroides spp.*, and *Clostridium spp.*, and host metabolic pathways (e.g., BCAAs, BA, and FFA) (Figs. 9B and 10B).

The microbiota composition and host metabolism were affected by CIH- and CSF-induced changes in body weight, fasting BG levels, and the weights of WAT, BAT, and liver. These results suggested that the structure of various fecal microorganisms was influenced by CIH and CSF, which was closely related to host metabolism and hepatic and adipose morphology.

## Discussion

In this study, we analyzed OSA-associated pathological changes in intestinal bacteria and metabolites, as well as their correlations with hepatic and adipose tissue morphology and lipid metabolism disorders. We found that CIH and CSF, with or without HFD-induced dysbiosis of intestinal bacteria and metabolites thereof, may independently or synergistically affect the risk of lipid-related metabolic complications. Adipose tissue and hepatic lipid metabolism involves a dynamic equilibrium between lipogenesis and lipolysis that maintains lipid homeostasis (17). C57BL/6 mice were the optimal model animal for simulating human lipid metabolism disorders (18). Abnormal lipid metabolism is suggested by body weight changes, as well as changes in the weight of subcutaneous or visceral WAT, visceral BAT and liver tissues, and an abnormal distribution of lipid droplets.

We found that CIH and CSF were associated with weight loss in murine models at the early stage of the experiment, even after HFD supplementation (Fig. 1B, D), which may correlate with a CIH- and CSF-induced reduction in the abundance of *A. muciniphila* (Figs. 4D, 5D, 10A and 10B), and a CSF-induced reduction in the abundance of *M. schaedleri* (Fig. 5D, 10B). Reduced abundances of the abovementioned bacteria led to decreased density of enteric goblet cells, increased intestinal permeability, impaired digestive function (19) and resulted in weight loss (Figs. 10A, 10B). The damaged intestinal barrier allows intestinal microbiome-derived components and metabolites to reach target organs, such as adipose and liver tissues.

Besides CIH- or CSF-related functional changes in intestinal bacteria promoted inhibition of the citrate cycle (Figs. 6K, 7K, 8B and 9B), a significant quantity of BAT was consumed to meet the energy requirement (Fig. 10A, B) (20). Additionally, BAT cells contained multiple lipid droplets and abundant mitochondria. Upregulation of mitochondrial uncoupling protein 1 (UCP1) activated BAT-mediated decomposition of triglycerides (TGs) into FFAs (21), producing energy for non-shivering heat generation through β-oxidation (22). CIH and CSF upregulated Firmicutes (Figs. 4C and 5C), and CIH significantly upregulated *Bifidobacterium spp.* and *Lactobacillus spp.* (Figs. 4D and 10A), which may act as strong peroxisome proliferator-activated receptor-γ (PPAR-γ) activators (23). PPAR-γ activation may upregulate UCP1 expression to promote BAT consumption (24), whereas increase BCAA degradation (Figs. 6H, 7H, 8A and 9A), which in turn affects body mass (Fig. 10A, B) (25). Similarly, weight loss decreased during the posterior experiments (Fig. 1B, D). Moreover, the CIH+HFD group had an increased abundance of Actinobacteria (Fig. 4C), which can synthesize PUFAs (26) from HFD and fatty acid derivatives (Fig. 8A), thereby activating PPAR-γ (27) to improve BAT metabolism in obese mice (Fig. 10A).

However, CIH and CSF intervention reduced the abundance of *Clostridium spp.* (Figs. 4D, 8B, 5D and 9B), leading to the inhibition of tryptophan absorption (Figs. 6I, 7I, 8A and 9A). *Clostridium spp.* converts raw tryptophan into various indole metabolites, such as 5-hydroxytryptamine (5-HT) and melatonin (28). The indole derivatives not only activate PPAR-γ to promote fatty acid thermogenic energy metabolism in rodent models (29), but also serve as an agonist to the aryl hydrocarbon receptor (AHR) to improve intestinal barrier function (30). For instance, melatonin may normalize the F/B ratio and abundance of *A. mucinphila* (29) to improve microbiome dysbiosis, in addition to regulating the circadian rhythm. Decreased tryptophan-derived indoles due to CIH- or CSF-related intestinal dysbiosis mediate intestinal permeability and BAT activation (Fig. 10A, B).

Excessive energy intake in our HFD group increased the body weight and caused obesity. HFD and CIH had a significant effect on BAT metabolism. Hypoxia was seen in obese mice (31); after CIH, excessive oxygen consumption during the β-oxidation process was blocked and FFAs were decomposed by BAT (32). We observed changes in the levels of FFAs under CIH and CSF conditions, which were accentuated by HFD supplementation (Figs. 6L and 7L). Furthermore, *Bifidobacterium spp.* and *Lactobacillus spp.* (Fig. 8A, B) were upregulated by CIH. *Clostridium spp.* (Figs. 8A, B and 9A, B) was downregulated by CIH and CSF (with or without HFD), resulting in excessive accumulation of FFAs due to altered dietary lipid absorption.

FFAs act as ligands to activate the Toll-like receptor (TLR) 2 pathway (33). FFAs may promote lipid metabolic instability and a pro-inflammatory state. Moreover, CIH, CSF, and HFD increased the levels of the pro-inflammatory bacterial compound lipopolysaccharide (LPS, derived from Gram-negative bacterial membranes), which serves as a ligand to activate TLR4 (34). The TLR family of pattern recognition receptors are associated with innate immunity, inflammation, and apoptosis of adipose tissue (34, 35). Activated TLRs inhibited β-oxidation and significantly increased the expression of pro-inflammatory factors such as monocyte chemoattractant protein-1 (MCP-1), interleukin-6 (IL-6), and tumor necrosis factor-alpha (TNF-α) (36), and the pro-apoptosis protein Bax (37). These autophagy-promoting signals inactivate the mitochondria and lipophagy in BAT (32), consistent with FFA-induced effects on autophagy (Fig. 10A, B) such as adipose fibrosis and fragmentation of lipid droplets in BAT due to CIH, CSF, and/or HFD (Figs. 2H and 3H). BAT inactivation (36) is eventually transformed into a crucial inducer of damage to lipid homeostasis.

In contrast to BAT, WAT cells contain only one lipid droplet and few mitochondria. Importantly, we did not observe WAT proliferation in response to CIH or CSF exposure alone. However, CIH or CSF combined with HFD led to an apparent increase in lipid deposition in IWAT and PWAT. CIH and CSF increased F/B ratio (Figs. 4C and 5C), consistent with the upregulation of glycerolipid metabolism (Figs. 6M and 7M), which promoted the increased TG deposition seen in WAT (38). Importantly, CSF upregulated the probiotic Oscillibacter spp. (Fig. 5D), which improves lipid metabolism (including of FFAs and glycerolipids) (Fig 9B) and inhibits pathological lipid deposition in IWAT and PWAT (Fig. 10B) (39).

Additionally, a large number of WAT cells were transformed into large vacuolated adipose cells (Figs. 2D, F and 3F). Furthermore, previous comparative studies of the expression profiles (40) demonstrated that small-sized adipocytes showed high expression levels of anti-inflammatory factors, such as adiponectin and resistin; conversely, large adipocytes that are insensitive to insulin highly expressed inflammatory factors, such as TNF-α and IL-6. In other words, CIH and CSF+HFD increased the number of pro-inflammatory WAT cells, which may contribute to subsequent disturbance of glucose (Fig. 1C, E). Therefore, apoptosis signal-regulating kinase1 may be specifically activated by TNF-α (41) or enterobacterial disorders (42). The CIH+HFD group demonstrated abnormal morphology and thin cell walls in the IWAT due to the pro-inflammatory and pro-apoptotic conditions (Fig. 2D).

Additionally, CIH intervention alone (Figs. 4C, D) significantly inhibited the probiotics Bacteroidetes and *Faecalibacterium prausnitzii*, and upregulated Actinobacteria and Desulfovibrio, leading to pro-inflammatory intestinal dysbiosis (43). After HFD supplementation, CSF decreased the abundance of Bacteroidetes and increased that of Proteobacteria (Fig. 5C), leading to pro-inflammatory intestinal microbiota. Combined with increased intestinal permeability, LPS promoted the secretion of MCP-1 from large adipocytes, resulting in macrophage accumulation and conversion into pro-inflammatory M1-type adipocyte macrophages (ATM) in WAT (44). The aggregation of M1-type ATM promoted the inflammatory cascade and C-reactive protein synthesis in the liver (45, 46). Therefore, WAT promoted inflammation, IR, and hepatic lipid deposition on exposure to OSA-related factors.

CIH and CSF-derived excessive ROS production caused oxidative damage to cellular macromolecules and altered the physical and chemical properties of liver membranes. Repeated oxidative stress depleted reduced glutathione (47), an essential free radical scavenger. CIH upregulated *Enterorhabdus spp.* and CSF down-regulated *Bacteroides spp.* (Figs. 4D, 5D, 8B and 9B), which inhibited glutamate absorption (with no significant change in glycine metabolism) and reduced glutathione synthesis (Figs. 6D, 7D, 6F, 7F, 8A and 9A). These changes significantly reduced the capacity for free radical elimination (48), leading to lysosome and mitochondria damage. HFD supplementation worsened the imbalance in the redox state, thereby leading to lipid peroxidation and subsequent liver steatosis.

Oil red staining demonstrated that CIH and CSF induced micro-vesicular lipid droplet deposition in hepatocytes, whereas HFD led to diffuse macro-vesicular hepatic steatosis (Figs. 2B and 3B), suggesting that OSA-related changes and dietary intervention had different and independent impacts on steatosis. CIH remarkably increased the F/B ratio (Fig 4C) (8), while CSF significantly inhibited polysaccharide fermentation (Figs. 6J, 7J, 8A and 9A) by decreasing Bacteroides (Figs. 5D and 9B), which decreases SCFAs synthesis (49). SCFAs protect the liver by upregulating the AMP protein kinase signaling pathway to activate PPAR-α in hepatic adipose tissue (50). Inhibition of the protective PPAR-α in hepatocytes due to insufficient SCFA synthesis, caused by CIH- or CSF-induced changes in intestinal bacteria and derived metabolites, is the root cause of liver lipid metabolism dyshomeostasis (Fig. 10A, B) (51).

Both CIH and CSF activated the sympathetic adrenal medulla system to release catecholamines, thereby mobilizing glucose and FFAs (52). However, HFD-induced obesity inhibited sympathetic excitability and induced the formation of lipid droplets in hepaticytes (53). CIH-, CSF-, and HFD-mediated intestinal bacteria dysbiosis resulted in stockpiling FFAs (Figs. 6L, 7L, 8A, B, and 9A, B), thereby exceeding the lipid acidification capacity of mitochondria. This leads to the deposition of FFAs in the liver through the portal vein (54). CIH stimulation promoted the expression of hypoxia-inducible factors-1α (55), and HFD upregulated the levels of stearyl CoA desaturase-1, fatty acid synthase (56), and genes involved in adipogenesis and lipid droplet deposition. Therefore, CIH, CSF, and HFD promoted the use of FFAs as raw materials to synthesize TGs and cholesterol (Fig. 10A, B), leading to hepatocyte steatosis (Figs. 2B and 3B) in non-alcoholic fatty liver disease (56, 57).

Even under physiological conditions, the liver consumes a large quantity of oxygen to convert excess dietary carbohydrates into TGs (57). The dual blood supply of the liver leads to a low physiological oxygen partial pressure, which makes hepatocytes, especially those around the central lobule, extremely sensitive to hypoxia. The morphological examination conducted in this study demonstrated that the hepatic vascular sinus region was the first region affected by CIH (and the most severely affected) (Fig. 2B).

The liver receives most of its blood supply directly from the intestine, which carries the intestinal flora through the portal vein to the liver. Therefore, the host intestinal microbiome and liver are intrinsically linked (58), such that dysbiosis of the intestinal bacterial and derived metabolites due to CIH and CSF leads to hepatic dysfunction (59). CIH, especially CIH+HFD, significantly upregulated *Bifidobacterium spp.* (Fig. 4D), which produces endogenous ethanol via fermentation to stimulate the oxidative stress in hepatic cells and exacerbates liver inflammation (60). Moreover, bacterial ethanol activated the TLRs (34) in hepatocytes to promote chronic hepatic inflammation due to increased intestinal permeability, and also raised the LPS level (61), consistent with the hepatic morphological changes seen in the CIH+HFD group (i.e., diffuse hepatic steatosis, cord disorders, and fibrosis) (Fig. 2A, B). CIH- and CSF-related bacterial dysbiosis upregulated LPS in the liver and affected peripheral lipid metabolism (62). Even low-dose LPS increased the synthesis of very-low-density lipoprotein in the liver, and high-dose LPS reduced lipoprotein decomposition and promoted dyslipidemia.

CIH and CSF were associated with increased intestinal permeability and levels of intestinal bacterial TLR ligands, which caused adipose and hepatic tissue inflammation and dyslipidosis (63). In OSA, adipose tissue may be the initial site of metabolic inflammation. Massive release of pro-inflammatory cytokines and activation of M1-type ATMs are the first metabolic derangements seen in OSA, followed by the gradual development of metabolic liver inflammation (64, 65). Considering the complex interactions in the CIH- and CSF-related intestinal microbiota-metabolome-adipose/liver axis, it is hypothesized that once hepatic metabolic inflammation is induced, the action of the liver enhances lipid metabolism disorders.

Dyshomeostasis of lipid and glucose metabolism often occur together. CIH and CSF activated the hypothalamic-pituitary-adrenal (HPA) cortex (66), leading to the release of catecholamines, glucocorticoids, and cortisol (Figs. 6Q and 7Q) and an increase in the level of fasting BG (Figs. 1C, E and 10A, B). Both CIH and CSF inhibited tryptophan absorption and indoles synthesis, which act as AHR agonists to promote secretion of the intestinal hormone GLP-1 and maintain glucose homeostasis (Fig. 10A, B) (30). Dysbiosis of the intestinal bacteria due to CIH or CSF alone downregulated the production of primary BAs (Figs. 6P, 7P, 8A and 9A). The abundances of Bacteroidetes and *Clostridium spp.*, which reduced the production of secondary BAs through the 7α-decarboxylation reaction process (67), was down-regulated by CIH and CSF (Figs. 4C, 4D, 5C, 5D). Secondary BAs act through Takeda G protein-coupled receptor 5 and promote GLP-1 release from the intestinal L cells (68). Therefore, insufficient secretion of endogenous GLP-1 due to CIH- and CSF-related changes in the intestinal bacteria and derived metabolites could disrupt glucose homeostasis. Additionally, decreased synthesis and increased consumption of BCAAs derived from the intestinal bacteria in CIH and CSF mice (Figs. 6H, 7H, 8A, 9A and 10A, B) positively correlated with IR (69). The CIH-, CSF-, and HFD-related gut microbiota dysbiosis resulted in very high FFAs levels (Figs. 6L, 7L, 8A and 9A), which activated the c-Jun N-terminal kinase and NF-κB pathways, leading to endoplasmic reticulum stress (70). In turn, this significantly affected the insulin sensitivity of adipocytes, causing glucose dyshomeostasis in adipocytes and aberrant BG levels (Fig. 10A, B).

In addition to the aforementioned metabolic disorders, CIH- and CSF-induced alterations of the gut microbiota, which predisposed the mice to CVDs, NDDs, and IDs (Fig. 8A, 9A). CIH induced periodic excitation of sympathetic and parasympathetic nerves, autonomic nervous dysfunction (71). Additionally, CIH-induced dysbiosis of the intestinal bacteria increased the homocysteine level (Figs. 6E and 8A), leading to endothelial damage through free oxygen radicals and nitric oxide, which caused vascular remodeling and lipid accumulation in the vascular wall (72). These changes were closely related to the development of CVD. CSF activated the HPA axis and neurons, resulting in excitatory toxicity and neuronal injury. Meanwhile, the increase abundance of Proteobacteria and the decrease abundance of Bacteroidetes due to CSF and CSF+HFD (Fig. 5D) decreased the levels of sphingolipid (Figs. 7R, 9A) and brain-derived neurotrophic factor through the enterobacteria-brain axis (73), which plays a crucial role in neuron survival, differentiation, and growth. Additionally, CSF down-regulated *E. coprostanoligenes* and endogenous vitamin B (Figs. 5D, 7S, 9B), resulting in decreased immunity (74) and an increased risk of IDs (Fig. 9B).

In conclusion, CIH and CSF regulated the levels of intestinal microbes (such as *A. mucinphila, Clostridium spp., Lactococcus spp.,* and *Bifidobacterium spp.*) through changes in many functional metabolites, such as tryptophan, FFAs, BCAA and Bas. These can affect adipose and hepatic tissue lipid metabolism and deposition (Table 1.). The CIH-gut microbiota-metabolite-adipose axis promoted metabolic inflammation, whereas the CIH-gut microbiota-metabolite-liver axis promoted hepatic steatosis. The CSF-gut microbiota-metabolite-adipose axis is crucial in the regulation of BAT-mediated energy metabolism.

**Table 1.**
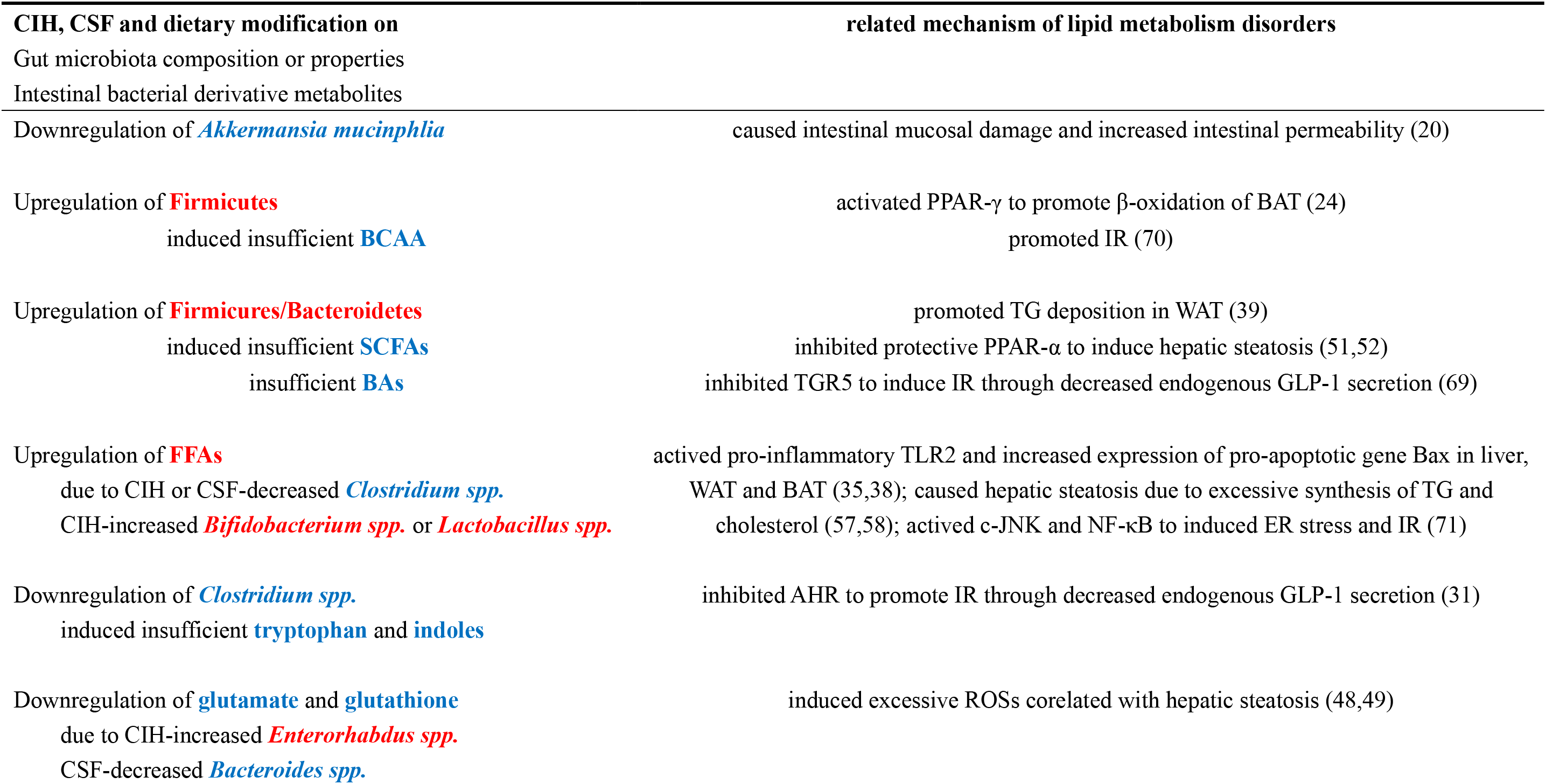
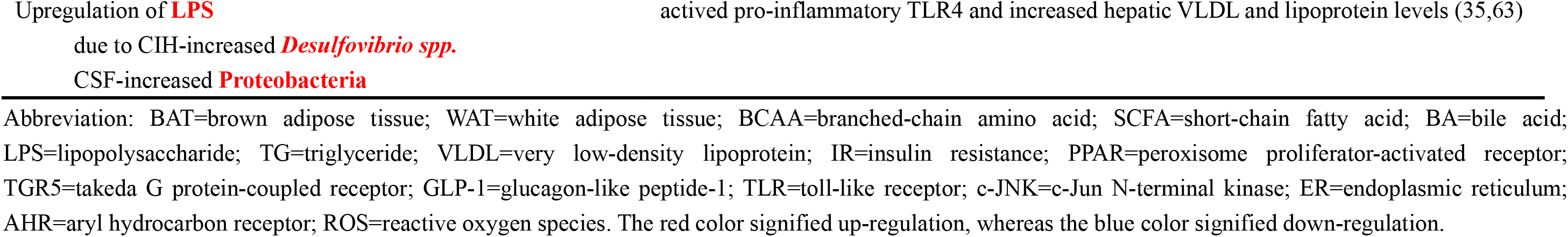
Commonalities between CIH and CSF (with or without HFD-fed) related modification on lipid metabolism through gut microbiota-metabolite-liver/adipose axis.

Our study had several limitations. First, due to the interaction and synergistic regulation of multiple functional metabolites, it was difficult to identify the key metabolites regulating changes in other metabolites. Third, functional metabolites with physiological and pathological effects were not comprehensively evaluated, and many metabolites may have been missed.

Gut microbes and their downstream metabolites are important for host health, linking diet and environmental factors in that context. Regulating the changes in intestinal flora associated with CIH and CSF may constitute a new approach to the prevention and treatment of OSA-induced lipid metabolic disorders. However, this emerging area of research requires further data. Our future studies should use tools such as metabolite annotation and gene integration to correlate gene sequences with metabolomics data, to generate multi-layer datasets (mRNA transcription, protein translation, and epigenetic modification). The mechanism of regulation of lipid metabolism by the intestinal microbiota and functional metabolites should be explored in sterile animals, using engineered bacteria (metabolic enzyme knockout or overexpression) and artificial compounds (75).

## References

1. Lyons MM, Bhatt NY, Pack AI, Magalang UJ. 2020. Global burden of sleep-disordered breathing and its implications. Respirology 25:690–702.

2. Benjafield AV, Ayas NT, Eastwood PR, Heinzer R, Ip MSM, Morrell MJ, Nunez CM, Patel SR, Penzel T, Pépin JL, Peppard PE, Sinha S, Tufik S, Valentine K, Malhotra A. 2019. Estimation of the global prevalence and burden of obstructive sleep apnoea: a literature-based analysis. Lancet Respir Med 7:687–698.

3. Punjabi NM. 2008. The epidemiology of adult obstructive sleep apnea. Proc Am Thorac Soc 5:136–43.

4. Andrade AG, Bubu OM, Varga AW, Osorio RS. 2018. The Relationship between Obstructive Sleep Apnea and Alzheimer’s Disease. J Alzheimers Dis 64:S255–s270.

5. Muraki I, Wada H, Tanigawa T. 2018. Sleep apnea and type 2 diabetes. J Diabetes Investig 9:991–997.

6. Drager LF, Togeiro SM, Polotsky VY, Lorenzi-Filho G. 2013. Obstructive sleep apnea: a cardiometabolic risk in obesity and the metabolic syndrome. J Am Coll Cardiol 62:569–76.

7. Zhang X, Wang S, Xu H, Yi H, Guan J, Yin S. 2021. Metabolomics and microbiome profiling as biomarkers in obstructive sleep apnoea: a comprehensive review. Eur Respir Rev 30.

8. Moreno-Indias I, Torres M, Montserrat JM, Sanchez-Alcoholado L, Cardona F, Tinahones FJ, Gozal D, Poroyko VA, Navajas D, Queipo-Ortuño MI, Farré R. 2015. Intermittent hypoxia alters gut microbiota diversity in a mouse model of sleep apnoea. Eur Respir J 45:1055–65.

9. Tripathi A, Melnik AV, Xue J, Poulsen O, Meehan MJ, Humphrey G, Jiang L, Ackermann G, McDonald D, Zhou D, Knight R, Dorrestein PC, Haddad GG. 2018. Intermittent Hypoxia and Hypercapnia, a Hallmark of Obstructive Sleep Apnea, Alters the Gut Microbiome and Metabolome. mSystems 3.

10. Conotte S, Tassin A, Conotte R, Colet JM, Zouaoui Boudjeltia K, Legrand A. 2018. Metabonomic profiling of chronic intermittent hypoxia in a mouse model. Respir Physiol Neurobiol 256:157–173.

11. Poroyko VA, Carreras A, Khalyfa A, Khalyfa AA, Leone V, Peris E, Almendros I, Gileles-Hillel A, Qiao Z, Hubert N, Farré R, Chang EB, Gozal D. 2016. Chronic Sleep Disruption Alters Gut Microbiota, Induces Systemic and Adipose Tissue Inflammation and Insulin Resistance in Mice. Sci Rep 6:35405.

12. Yoon DW, Kwon HN, Jin X, Kim JK, Lee SK, Park S, Yun CH, Shin C. 2019. Untargeted metabolomics analysis of rat hippocampus subjected to sleep fragmentation. Brain Res Bull 153:74–83.

13. Wang Y, Lee MYK, Mak JCW, Ip MSM. 2019. Low-Frequency Intermittent Hypoxia Suppresses Subcutaneous Adipogenesis and Induces Macrophage Polarization in Lean Mice. Diabetes Metab J 43:659–674.

14. Li Y, Panossian LA, Zhang J, Zhu Y, Zhan G, Chou YT, Fenik P, Bhatnagar S, Piel DA, Beck SG, Veasey S. 2014. Effects of chronic sleep fragmentation on wake-active neurons and the hypercapnic arousal response. Sleep 37:51–64.

15. Dunn WB, Wilson ID, Nicholls AW, Broadhurst D. 2012. The importance of experimental design and QC samples in large-scale and MS-driven untargeted metabolomic studies of humans. Bioanalysis 4:2249–64.

16. Langille MG, Zaneveld J, Caporaso JG, McDonald D, Knights D, Reyes JA, Clemente JC, Burkepile DE, Vega Thurber RL, Knight R, Beiko RG, Huttenhower C. 2013. Predictive functional profiling of microbial communities using 16S rRNA marker gene sequences. Nat Biotechnol 31:814–21.

17. Olzmann JA, Carvalho P. 2019. Dynamics and functions of lipid droplets. Nat Rev Mol Cell Biol 20:137–155.

18. Collins S, Martin TL, Surwit RS, Robidoux J. 2004. Genetic vulnerability to diet-induced obesity in the C57BL/6J mouse: physiological and molecular characteristics. Physiol Behav 81:243–8.

19. Le Roy T, Llopis M, Lepage P, Bruneau A, Rabot S, Bevilacqua C, Martin P, Philippe C, Walker F, Bado A, Perlemuter G, Cassard-Doulcier AM, Gérard P. 2013. Intestinal microbiota determines development of non-alcoholic fatty liver disease in mice. Gut 62:1787–94.

20. Kang Y, Cai Y. 2018. The development of probiotics therapy to obesity: a therapy that has gained considerable momentum. Hormones (Athens) 17:141–151.

21. Bai Y, Meng L, Han L, Jia Y, Zhao Y, Gao H, Kang R, Wang X, Tang D, Dai E. 2019. Lipid storage and lipophagy regulates ferroptosis. Biochem Biophys Res Commun 508:997–1003.

22. Sidossis L, Kajimura S. 2015. Brown and beige fat in humans: thermogenic adipocytes that control energy and glucose homeostasis. J Clin Invest 125:478–86.

23. Nepelska M, de Wouters T, Jacouton E, Béguet-Crespel F, Lapaque N, Doré J, Arulampalam V, Blottière HM. 2017. Commensal gut bacteria modulate phosphorylation-dependent PPARγ transcriptional activity in human intestinal epithelial cells. Sci Rep 7:43199.

24. Moreno-Navarrete JM, Fernandez-Real JM. 2019. The gut microbiota modulates both browning of white adipose tissue and the activity of brown adipose tissue. Rev Endocr Metab Disord 20:387–397.

25. Blanchard PG, Moreira RJ, Castro É, Caron A, Côté M, Andrade ML, Oliveira TE, Ortiz-Silva M, Peixoto AS, Dias FA, Gélinas Y, Guerra-Sá R, Deshaies Y, Festuccia WT. 2018. PPARγ is a major regulator of branched-chain amino acid blood levels and catabolism in white and brown adipose tissues. Metabolism 89:27–38.

26. Lu J, Borthwick F, Hassanali Z, Wang Y, Mangat R, Ruth M, Shi D, Jaeschke A, Russell JC, Field CJ, Proctor SD, Vine DF. 2011. Chronic dietary n-3 PUFA intervention improves dyslipidaemia and subsequent cardiovascular complications in the JCR:LA-cp rat model of the metabolic syndrome. Br J Nutr 105:1572–82.

27. Marion-Letellier R, Savoye G, Ghosh S. 2016. Fatty acids, eicosanoids and PPAR gamma. Eur J Pharmacol 785:44–49.

28. Zelante T, Iannitti RG, Cunha C, De Luca A, Giovannini G, Pieraccini G, Zecchi R, D’Angelo C, Massi-Benedetti C, Fallarino F, Carvalho A, Puccetti P, Romani L. 2013. Tryptophan catabolites from microbiota engage aryl hydrocarbon receptor and balance mucosal reactivity via interleukin-22. Immunity 39:372–85.

29. Yin J, Li Y, Han H, Chen S, Gao J, Liu G, Wu X, Deng J, Yu Q, Huang X, Fang R, Li T, Reiter RJ, Zhang D, Zhu C, Zhu G, Ren W, Yin Y. 2018. Melatonin reprogramming of gut microbiota improves lipid dysmetabolism in high-fat diet-fed mice. J Pineal Res 65:e12524.

30. Xu P, Wang J, Hong F, Wang S, Jin X, Xue T, Jia L, Zhai Y. 2017. Melatonin prevents obesity through modulation of gut microbiota in mice. J Pineal Res 62.

31. Ye J. 2009. Emerging role of adipose tissue hypoxia in obesity and insulin resistance. Int J Obes (Lond) 33:54–66.

32. Singh R, Kaushik S, Wang Y, Xiang Y, Novak I, Komatsu M, Tanaka K, Cuervo AM, Czaja MJ. 2009. Autophagy regulates lipid metabolism. Nature 458:1131–5.

33. Mocanu AO, Mulya A, Huang H, Dan O, Shimizu H, Batayyah E, Brethauer SA, Dinischiotu A, Kirwan JP. 2015. Effect of Roux-en-Y Gastric Bypass on the NLRP3 Inflammasome in Adipose Tissue from Obese Rats. PLoS One 10:e0139764.

34. Satoh T, Akira S. 2016. Toll-Like Receptor Signaling and Its Inducible Proteins. Microbiol Spectr 4.

35. Bastard JP, Maachi M, Lagathu C, Kim MJ, Caron M, Vidal H, Capeau J, Feve B. 2006. Recent advances in the relationship between obesity, inflammation, and insulin resistance. Eur Cytokine Netw 17:4–12.

36. Li Y, Wang C, Lu J, Huang K, Han Y, Chen J, Yang Y, Liu B. 2019. PPAR δ inhibition protects against palmitic acid-LPS induced lipidosis and injury in cultured hepatocyte L02 cell. Int J Med Sci 16:1593–1603.

37. Calderon-Dominguez M, Mir JF, Fucho R, Weber M, Serra D, Herrero L. 2016. Fatty acid metabolism and the basis of brown adipose tissue function. Adipocyte 5:98–118.

38. Bo TB, Wen J, Zhao YC, Tian SJ, Zhang XY, Wang DH. 2020. Bifidobacterium pseudolongum reduces triglycerides by modulating gut microbiota in mice fed high-fat food. J Steroid Biochem Mol Biol 198:105602.

39. Yuan Z, Yan W, Wen C, Zheng J, Yang N, Sun C. 2020. Enterotype identification and its influence on regulating the duodenum metabolism in chickens. Poult Sci 99:1515–1527.

40. Weyer C, Foley JE, Bogardus C, Tataranni PA, Pratley RE. 2000. Enlarged subcutaneous abdominal adipocyte size, but not obesity itself, predicts type II diabetes independent of insulin resistance. Diabetologia 43:1498–506.

41. Coppack SW. 2001. Pro-inflammatory cytokines and adipose tissue. Proc Nutr Soc 60:349–56.

42. Lucchini FC, Wueest S, Challa TD, Item F, Modica S, Borsigova M, Haim Y, Wolfrum C, Rudich A, Konrad D. 2020. ASK1 inhibits browning of white adipose tissue in obesity. Nat Commun 11:1642.

43. Zhu L, Baker SS, Gill C, Liu W, Alkhouri R, Baker RD, Gill SR. 2013. Characterization of gut microbiomes in nonalcoholic steatohepatitis (NASH) patients: a connection between endogenous alcohol and NASH. Hepatology 57:601–9.

44. Engin AB. 2017. Adipocyte-Macrophage Cross-Talk in Obesity. Adv Exp Med Biol 960:327–343.

45. Gao Z, Zhang J, Henagan TM, Lee JH, Ye X, Wang H, Ye J. 2015. P65 inactivation in adipocytes and macrophages attenuates adipose inflammatory response in lean but not in obese mice. Am J Physiol Endocrinol Metab 308:E496–505.

46. Schmidt-Arras D, Rose-John S. 2016. IL-6 pathway in the liver: From physiopathology to therapy. J Hepatol 64:1403–15.

47. Paamoni-Keren O, Silberstein T, Burg A, Raz I, Mazor M, Saphier O, Weintraub AY. 2007. Oxidative stress as determined by glutathione (GSH) concentrations in venous cord blood in elective cesarean delivery versus uncomplicated vaginal delivery. Arch Gynecol Obstet 276:43–6.

48. Mardinoglu A, Shoaie S, Bergentall M, Ghaffari P, Zhang C, Larsson E, Bäckhed F, Nielsen J. 2015. The gut microbiota modulates host amino acid and glutathione metabolism in mice. Mol Syst Biol 11:834.

49. Zhao Y, Wu J, Li JV, Zhou NY, Tang H, Wang Y. 2013. Gut microbiota composition modifies fecal metabolic profiles in mice. J Proteome Res 12:2987–99.

50. Kersten S, Stienstra R. 2017. The role and regulation of the peroxisome proliferator activated receptor alpha in human liver. Biochimie 136:75–84.

51. Montagner A, Polizzi A, Fouché E, Ducheix S, Lippi Y, Lasserre F, Barquissau V, Régnier M, Lukowicz C, Benhamed F, Iroz A, Bertrand-Michel J, Al Saati T, Cano P, Mselli-Lakhal L, Mithieux G, Rajas F, Lagarrigue S, Pineau T, Loiseau N, Postic C, Langin D, Wahli W, Guillou H. 2016. Liver PPARα is crucial for whole-body fatty acid homeostasis and is protective against NAFLD. Gut 65:1202–14.

52. Verberne AJ, Korim WS, Sabetghadam A, Llewellyn-Smith IJ. 2016. Adrenaline: insights into its metabolic roles in hypoglycaemia and diabetes. Br J Pharmacol 173:1425–37.

53. Troisi RJ, Weiss ST, Parker DR, Sparrow D, Young JB, Landsberg L. 1991. Relation of obesity and diet to sympathetic nervous system activity. Hypertension 17:669–77.

54. Drager LF, Li J, Reinke C, Bevans-Fonti S, Jun JC, Polotsky VY. 2011. Intermittent hypoxia exacerbates metabolic effects of diet-induced obesity. Obesity (Silver Spring) 19:2167–74.

55. Li J, Bosch-Marce M, Nanayakkara A, Savransky V, Fried SK, Semenza GL, Polotsky VY. 2006. Altered metabolic responses to intermittent hypoxia in mice with partial deficiency of hypoxia-inducible factor-1alpha. Physiol Genomics 25:450–7.

56. Shin MR, Shin SH, Roh SS. 2020. Diospyros kaki and Citrus unshiu Mixture Improves Disorders of Lipid Metabolism in Nonalcoholic Fatty Liver Disease. Can J Gastroenterol Hepatol 2020:8812634.

57. Zechner R, Kienesberger PC, Haemmerle G, Zimmermann R, Lass A. 2009. Adipose triglyceride lipase and the lipolytic catabolism of cellular fat stores. J Lipid Res 50:3–21.

58. Woodhouse CA, Patel VC, Singanayagam A, Shawcross DL. 2018. Review article: the gut microbiome as a therapeutic target in the pathogenesis and treatment of chronic liver disease. Aliment Pharmacol Ther 47:192–202.

59. Aqel B, DiBaise JK. 2015. Role of the Gut Microbiome in Nonalcoholic Fatty Liver Disease. Nutr Clin Pract 30:780–6.

60. Cope K, Risby T, Diehl AM. 2000. Increased gastrointestinal ethanol production in obese mice: implications for fatty liver disease pathogenesis. Gastroenterology 119:1340–7.

61. Parlesak A, Schäfer C, Schütz T, Bode JC, Bode C. 2000. Increased intestinal permeability to macromolecules and endotoxemia in patients with chronic alcohol abuse in different stages of alcohol-induced liver disease. J Hepatol 32:742–7.

62. Read TE, Harris HW, Grunfeld C, Feingold KR, Kane JP, Rapp JH. 1993. The protective effect of serum lipoproteins against bacterial lipopolysaccharide. Eur Heart J 14 Suppl K:125–9.

63. Csak T, Ganz M, Pespisa J, Kodys K, Dolganiuc A, Szabo G. 2011. Fatty acid and endotoxin activate inflammasomes in mouse hepatocytes that release danger signals to stimulate immune cells. Hepatology 54:133–44.

64. Saltzman ET, Palacios T, Thomsen M, Vitetta L. 2018. Intestinal Microbiome Shifts, Dysbiosis, Inflammation, and Non-alcoholic Fatty Liver Disease. Front Microbiol 9:61.

65. Wright KP, Jr., Drake AL, Frey DJ, Fleshner M, Desouza CA, Gronfier C, Czeisler CA. 2015. Influence of sleep deprivation and circadian misalignment on cortisol, inflammatory markers, and cytokine balance. Brain Behav Immun 47:24–34.

66. Vargas I, Lopez-Duran N. 2017. Investigating the effect of acute sleep deprivation on hypothalamic-pituitary-adrenal-axis response to a psychosocial stressor. Psychoneuroendocrinology 79:1–8.

67. Ridlon JM, Hylemon PB. 2012. Identification and characterization of two bile acid coenzyme A transferases from Clostridium scindens, a bile acid 7α-dehydroxylating intestinal bacterium. J Lipid Res 53:66–76.

68. Ronveaux CC, Tomé D, Raybould HE. 2015. Glucagon-like peptide 1 interacts with ghrelin and leptin to regulate glucose metabolism and food intake through vagal afferent neuron signaling. J Nutr 145:672–80.

69. Zhou M, Shao J, Wu CY, Shu L, Dong W, Liu Y, Chen M, Wynn RM, Wang J, Wang J, Gui WJ, Qi X, Lusis AJ, Li Z, Wang W, Ning G, Yang X, Chuang DT, Wang Y, Sun H. 2019. Targeting BCAA Catabolism to Treat Obesity-Associated Insulin Resistance. Diabetes 68:1730–1746.

70. Wenfeng Z, Yakun W, Di M, Jianping G, Chuanxin W, Chun H. 2014. Kupffer cells: increasingly significant role in nonalcoholic fatty liver disease. Ann Hepatol 13:489–95.

71. Baguet JP, Barone-Rochette G, Tamisier R, Levy P, Pépin JL. 2012. Mechanisms of cardiac dysfunction in obstructive sleep apnea. Nat Rev Cardiol 9:679–88.

72. Koller A, Szenasi A, Dornyei G, Kovacs N, Lelbach A, Kovacs I. 2018. Coronary Microvascular and Cardiac Dysfunction Due to Homocysteine Pathometabolism; A Complex Therapeutic Design. Curr Pharm Des 24:2911–2920.

73. Schmitt K, Holsboer-Trachsler E, Eckert A. 2016. BDNF in sleep, insomnia, and sleep deprivation. Ann Med 48:42–51.

74. Spinas E, Saggini A, Kritas SK, Cerulli G, Caraffa A, Antinolfi P, Pantalone A, Frydas A, Tei M, Speziali A, Saggini R, Pandolfi F, Conti P. 2015. CROSSTALK BETWEEN VITAMIN B AND IMMUNITY. J Biol Regul Homeost Agents 29:283–8.

75. Finegold SM, Dowd SE, Gontcharova V, Liu C, Henley KE, Wolcott RD, Youn E, Summanen PH, Granpeesheh D, Dixon D, Liu M, Molitoris DR, Green JA, 3rd. 2010. Pyrosequencing study of fecal microflora of autistic and control children. Anaerobe 16:444–53.

